# Recovery of the gut microbiome following enteric infection and persistence of antimicrobial resistance genes in specific microbial hosts

**DOI:** 10.1101/2023.01.13.523990

**Authors:** Zoe A. Hansen, Karla A. Vasco, James T. Rudrik, Kim T. Scribner, Lixin Zhang, Shannon D. Manning

## Abstract

Enteric pathogens cause widespread foodborne illness and are increasingly found to harbor antimicrobial resistance. The ecological impact of these pathogens on the human gut microbiome and resistome, however, has yet to be fully elucidated. This study applied shotgun metagenome sequencing to stools from 60 patients (cases) with enteric bacterial infections for comparison to stools collected from the same patients’ post-recovery (follow-ups). Overall, the case samples harbored more antimicrobial resistance genes (ARGs) and had greater resistome diversity than the follow-up samples (p<0.001), while follow-ups had much more diverse microbiomes (p<0.001). Although cases were primarily defined by genera *Escherichia, Salmonella*, and *Shigella* along with ARGs for multi-compound and multidrug resistance, follow-ups had a greater abundance of Bacteroidetes and Firmicutes phyla and genes for tetracycline, macrolides, lincosamides, and streptogramins (MLS), and aminoglycoside resistance. A host-tracking analysis revealed that *Escherichia* was the primary carrier of ARGs in both cases and follow-ups, with a greater abundance occurring during infection. Eleven distinct extended spectrum beta-lactamases (ESBLs) were identified during infection, some of which appear to be lost or transferred to different microbial hosts upon recovery. The increasing incidence of disease caused by foodborne pathogens, coupled with their evolving role in harboring and transferring antimicrobial resistance determinants within communities, justifies further examination of the repercussions of enteric infection on human gut ecology.

## Introduction

Foodborne illness caused by enteric pathogens impacts ~9.4 million people in the United States each year, with over one-third being attributed to bacterial pathogens [1]. In 2019, the Centers for Disease Control and Prevention (CDC) documented a marked increase in the incidence of foodborne infection caused by *Campylobacter* and Shiga toxin-producing *Escherichia coli* (STEC) [2]. *Salmonella* and *Shigella* also contribute to a high incidence of infections, though case numbers remained unchanged relative to previous years. In addition to their role in enteric disease, *Campylobacter*, non-Typhoidal *Salmonella*, *Shigella*, and members of *Enterobacteriaceae* (e.g., *Escherichia*) have been classified by the CDC as serious threats for harboring and transmitting antimicrobial resistance [2]. Indeed, each of these pathogens have been shown to transfer ARGs horizontally within and between microbial species residing in a niche [3]. Such resistance determinants can cross environmental boundaries, thereby increasing frequencies within different hosts and environments and enhancing the likelihood of horizontal gene transfer (HGT).

The consequences of enteric infection on the health of the human gut microbiome are not fully understood. Prior studies conducted in our lab showed a marked decrease in gut microbiota diversity attributed to enteric infection [4]. This lack of diversity was suggested to reduce beneficial microbially-mediated metabolism and exacerbate gut inflammation [5]. Others have also demonstrated an increase in the proportion of Proteobacteria upon infection with *Salmonella*, *Campylobacter*, *Shigella*, and other pathogens in multiple host organisms [6–9]. More recently, we have documented shifts in the gut resistome, or compilation of antimicrobial resistance genes (ARGs), among patients with *Campylobacter* infections when compared to their healthy family members [10]. The potential ecological repercussions relevant to recovery from enteric infection, however, have yet to be explored. If the microbiome demonstrates a certain degree of resilience, then perturbations should not be felt with such amplitude and be resolved over time [11]. In the context of pathogen invasion, various ecological interactions such as direct antagonism from commensal microbes, resource competition and competitive exclusion, and secondary metabolite production, must be considered [12, 13]. Each of these factors may influence the success of an enteric pathogen in the gut environment and the ability of the human host to recover from the acute infection.

Consideration must also be given to the invading pathogen, which can potentially introduce virulence and antimicrobial resistance determinants into the gut community. Indeed, pathogens harboring ARGs can transfer these to other gut microbes during infection or vice versa, thereby transforming the gut into a resistance gene reservoir [14]. This reservoir is particularly concerning given that pathobionts found in the community can acquire genetic factors that encode for pathogenic properties as well as resistance to clinically important antibiotics. Because infection with enteric pathogens was shown to alter the relative abundance of certain microbial populations in the gut [4], it is probable that ARGs harbored by microbes that “bloom” during infection will also increase in abundance. Although new sequence-based approaches have been developed to identify the microbial hosts of specific ARGs in different environments [15], these have not been applied to enteric infections.

Consequently, we used shotgun metagenome data to determine how infection by and recovery from enteric pathogens influences the human gut resistome and microbiome. We also sought to identify which microbial hosts harbor ARGs to advance understanding of how drug resistance spreads and is maintained within the dysbiotic and healthy gut microbiome. Further defining the impacts of these infections on the makeup and function of the gut microbiome is necessary to counteract the dissemination of drug resistance and discover novel therapeutic solutions.

## Methods

### Sample collection and sequencing

Sixty stools were obtained from patients with enteric infections (cases) caused by *Campylobacter* (n=24), *Salmonella* (n=29) *Shigella* (n=4) and Shiga toxin-producing *E. coli* (STEC) (n=3) from, 2011-2015. Stools were preserved in Cary-Blair transport media and submitted to the Michigan Department of Health and Human Services (MDHHS) in collaboration with four hospitals as described [4]. Patient demographics, exposures, and symptoms were reported through the Michigan Disease Surveillance System (MDSS). Counties were classified as ‘rural’ or ‘urban’ as was done in our prior analysis [10]. Each patient submitted a follow-up sample 1 week to 29 weeks after their acute infection, yielding 120 paired samples for analysis. Moreover, 91 household members (controls) linked to 38 of the 60 patients submitted stools for comparison 5-29 weeks after the cases’ infection. Resistome data from *Campylobacter* patients were examined previously [10], though no prior metagenome analyses were performed on the post-recovery samples.

Metagenomic DNA was extracted, sheared, and normalized as described [4]. Libraries were constructed using the TruSeq Nano library kit (Illumina, Inc., San Diego, CA, USA) and shotgun sequencing was performed in four runs using an Illumina HiSeq 2500. Reads were demultiplexed at the MSU Research Technology Support Facility and poor quality and contaminated samples were removed after filtering.

### Reads-based identification of antimicrobial resistance genes (ARGs)

The AmrPlusPlus v2.0 pipeline was used for quality control checking, aligning, and annotating metagenomic fragments with the MEGARes 2.0 database [16] and previously described parameters [10]. Reads were mapped to human genome GRCh38 (GRCh38_latest_genomic.fna.gz, downloaded December 2020) in RefSeq using the Burrows-Wheeler Aligner (BWA) [17] and removed using SAMTools [18] and BEDTools [19]. The non-host FASTQ files were stored and aligned to MEGARes 2.0 to identify ARGs using default values for the BWA and SAMTools. Reads were deduplicated and annotated with ResistomeAnalyzer (identity threshold of ≥80%) to quantify ARG abundance per sample, while RarefactionAnalzyer estimated sequencing depth. Single nucleotide polymorphisms (SNPs) requiring specific haplotypes to be classified as ARGs were also extracted for confirmation using the Resistance Gene Identifier via the Comprehensive Antibiotic Resistance Database [20]. Following annotation and quantifying ARG abundances, MicrobeCensus [21] was used to determine the average genome size and number of genome equivalents (GE) for normalizing ARG and taxonomic abundances (**Additional file 1**). Lastly, metagenomic coverage was estimated with Nonpareil [22] (**Additional file 2**).

### Assembly-based identification of ARGs

The non-host FASTQ files were used for metagenome assembly after employing BBTools for paired end read merging using the ‘bbmerge-auto.sh’ script. Reads that failed merging were error-corrected using Tadpole [23] and reexamined. If merging continued to fail, reads were extended 20 bp and merging was iterated up to five additional times or unmerged reads were included. Assembly was performed with MEGAHIT [24] using the merged and paired-end reads. The Quality Assessment Tool for Genome Assemblies [25] evaluated assembly quality and coverage (**Additional file 3**).

A custom workflow was developed using anvi’o to analyze microbial genomes from metagenomes as described [26]. Briefly, assembled contigs were reformatted using ‘anvi-script-reformat-fasta’ to generate a contigs database per sample with ‘anvi-gen-contigs-database’. The script ‘anvi-run-hmms’ was used to populate the contigs database with hits detected using Hidden Markov Models, which improves assembly annotation. Prodigal [27] was used in the script ‘anvi-get-sequences-for-gene-calls’ to obtain amino acid sequences of assembled genes for use in the ARG-carrying contigs (ACC) analysis.

### Classifying microbial taxa

Non-host paired-end reads were taxonomically annotated with Kaiju, a protein-based classifier that translates reads to amino acid sequences while searching for maximum exact matches (MEMs) among microbial reference genomes [28]. The National Center for Biotechnology Information (NCBI) BLAST *nr* reference database was used with previously published parameters [10]. Raw abundances of reads assigned to taxa were normalized by the estimated number of GEs. Those sequencing reads without enough resolution were categorized as “unassigned”, which comprised ≥50% of annotated reads at the genus and species levels. The composition analysis was restricted to assigned reads.

### Identifying bacterial hosts harboring ARGs

Gene calls from anvi’o were used to identify ARG-carrying contigs (ACCs) by aligning the amino acid sequences to the HMD-ARG database [29] using DIAMOND [30] using a modified pipeline that was described previously [15, 31]. The SAM files were filtered to identify contigs with gene hits, and Seqtk (https://github.com/lh3/seqtk) was used to extract the ACCs from the genes as a FASTA file for alignment to the BLAST database v5.0 using blastp. An E-value of 0.00001 cutoff was used with a maximum of 50 target sequences (i.e., 50 matches per contig). One *Campylobacter* sample could not be annotated and was excluded along with the paired follow-up sample leaving 59 pairs (118 samples) for analysis.

Alignment output was used to identify taxa associated with each ARG on a contig. Since 50 matches were allowed per contig, a custom Python script ‘ERIN_ACCpipeline_blastp_merge’ was used to quantify the average proportion of each genus per sample on the ACCs and the average percentage of different ARGs per genus within all ACCs in a sample. Taxa with the most hits per contig were considered the most likely to harbor a given ARG.

### Abundance and diversity analyses

The identity and diversity of ARGs and taxa were determined among all samples. For the resistome analyses, the gene, group, mechanism, class, and type levels were used [16]. Actual estimated abundance of ARGs and taxa was determined by normalizing raw abundance counts to the number of GE per sample. Relative abundance was calculated by dividing the number of GE-normalized reads assigned to a specific feature by the total number of GE-normalized reads for that sample. Alpha diversity metrics such as richness, Shannon diversity, and the Pielou’s evenness score were estimated using the vegan package [32] in R (https://www.R-project.org/). Nonparametric tests evaluated differences between groups and the Shapiro-Wilk test indicated that both the resistome and microbiome data were not normally distributed (**Additional file 4**).

The Wilcoxon signed-rank test was used to detect significant differences between paired samples, whereas the Wilcoxon rank-sum test was applied to unpaired samples. Beta diversity metrics and ordination plots (e.g., Principal Coordinate Analysis (PCoA)) based on Bray-Curtis dissimilarity at the gene and group (ARGs) or species and genus (taxa) levels were also estimated with vegan [32]. The overall mean dissimilarity among cases and follow-ups was compared to the mean dissimilarity between paired samples using a Welch’s t-test (**Additional file 5**). A Permutational Analysis of Variance (PERMANOVA) was calculated using the Bray-Curtis dissimilarities in R to assess differences in centroids (mean) between cases and follow-ups for both the resistome and microbiome composition; Permutational Analysis of Multivariate Dispersion (PERMDISP) detected differences in dispersion (degree of spread) of these groups.

### Differential abundance of taxa and ARGs

To assess representative features in cases and follow-ups, MMUPHin was used to construct general linear models relating sample features to relative abundances [33]. Batch adjustment of relative abundance data was performed by sequencing run, which significantly influenced the distribution of points in the microbiome ordination (**Additional file 6**). To identify differentially abundant ARGs and taxa, a linear model was constructed with follow-ups serving as the reference for the fixed effect. Age in years, average genome size, number of GE, year of collection, and use of antibiotics were included as covariates. Significance values were adjusted using the Benjamini-Hochberg method of correction for multiple hypothesis testing (q-value representing False Discovery Rate). The Analysis of Compositions of Microbiomes with Bias Correction (ANCOM-BC) method [34], which considers absolute abundances from the GE-normalized counts as input but cannot implement a mixed model with fixed and random effects, was used for differential abundance testing. The data were concordant with MMUPHin data at each comparison level (**Additional file 7**), though differences in rank of correlation was observed for some features.

### Identification of continuous population structure

MMUPHin [33] was also used to identify continuous population structure from the microbiome and resistome abundance data to identify taxonomic or resistance gene tradeoffs that impact data structure in ordination. The ‘continous_discover()’ function was applied to the abundance data, which performs unsupervised continuous structure discovery using Principal Components Analysis (PCA). Continuous structure scores (called “loadings”) that comprise the top principal components were compared across batches to identify “consensus” loadings assigned to microbial features. The ‘var_perc_cutoff()’ parameter, which filters out the top components accounting for a set proportion of the variability within the samples, was set to 0.75 for phylum and ARG class levels, 0.50 for genus and ARG groups, and 0.40 for species. Plots were constructed to visualize the drivers of continuous data structure and to overlay data onto ordination plots based on Bray-Curtis dissimilarity of microbiome or resistome relative abundances.

## Results

### Study population

Among the 60 cases, 28 were male (46.7%) and 32 were female (53.3%) ranging between 1.5 and 90 years of age; most patients were between 19-64 years (n=26; 43.3%) or less than 9 (n=16; 26.7%). No difference in the proportion of stool submissions was observed by year, though the fewest (n=13.3%) were recovered in 2011 and the most (36.7%) in 2013. Among the 59 patients reporting symptoms, 50 (84.8%) had abdominal pain, 57 (96.6%) reported diarrhea, and 22 (37.3%) reported blood in the stool. Seventeen (28.3%) cases required hospitalization and 33 (55.0%) resided in a rural area. Most cases did not take antibiotics within two weeks of sampling, though two (3.3%) reported amoxicillin use, while five (8.3%) reported use of amoxicillin (n=2), azithromycin (n=1), ciprofloxacin (n=1), or an unknown antibiotic (n=1) before submitting the follow-up sample. Most follow-up samples were collected 51-100 days (n=20; 33.9%) or 101-150 days (n=28; 47.5%) post-infection, however, a small number was submitted ≤50 (n=4; 6.78%) or >150 (n=7;11.9%) days after the initial sample was collected; the date was missing for one patient. The range of follow-up submissions was 8 to 205 days post-recovery with an average of 107.9 days.

### Changes in resistome composition and diversity post-recovery

Among the 120 stool samples, 1,212 ARGs were identified encoding resistance to biocides, antibiotic drugs, metals, and multi-compound substrates comprising 474 distinct gene groups or operons. These genes represented 120 distinct mechanisms conferring resistance to 44 classes of compounds. In all, the case samples had significantly more diverse resistomes than follow-up samples with a greater mean ARG richness (S_cases_=254 vs. S_follow-ups_=103; p=4.5e-10) (**Figure 1**). The Shannon Diversity Index was also greater in cases than follow-ups (H_cases_=4.79 vs. H_follow-ups_=3.36; p=2.1e-10) as was the Pielou’s evenness index (J’_cases_=0.87 vs. J’_follow-ups_=0.80; p=8.1e-10). Notably, the family member controls did not significantly differ from follow-up samples, suggesting recovery to a “normal” state post-infection (**Additional file 8**).

**Figure 1.**
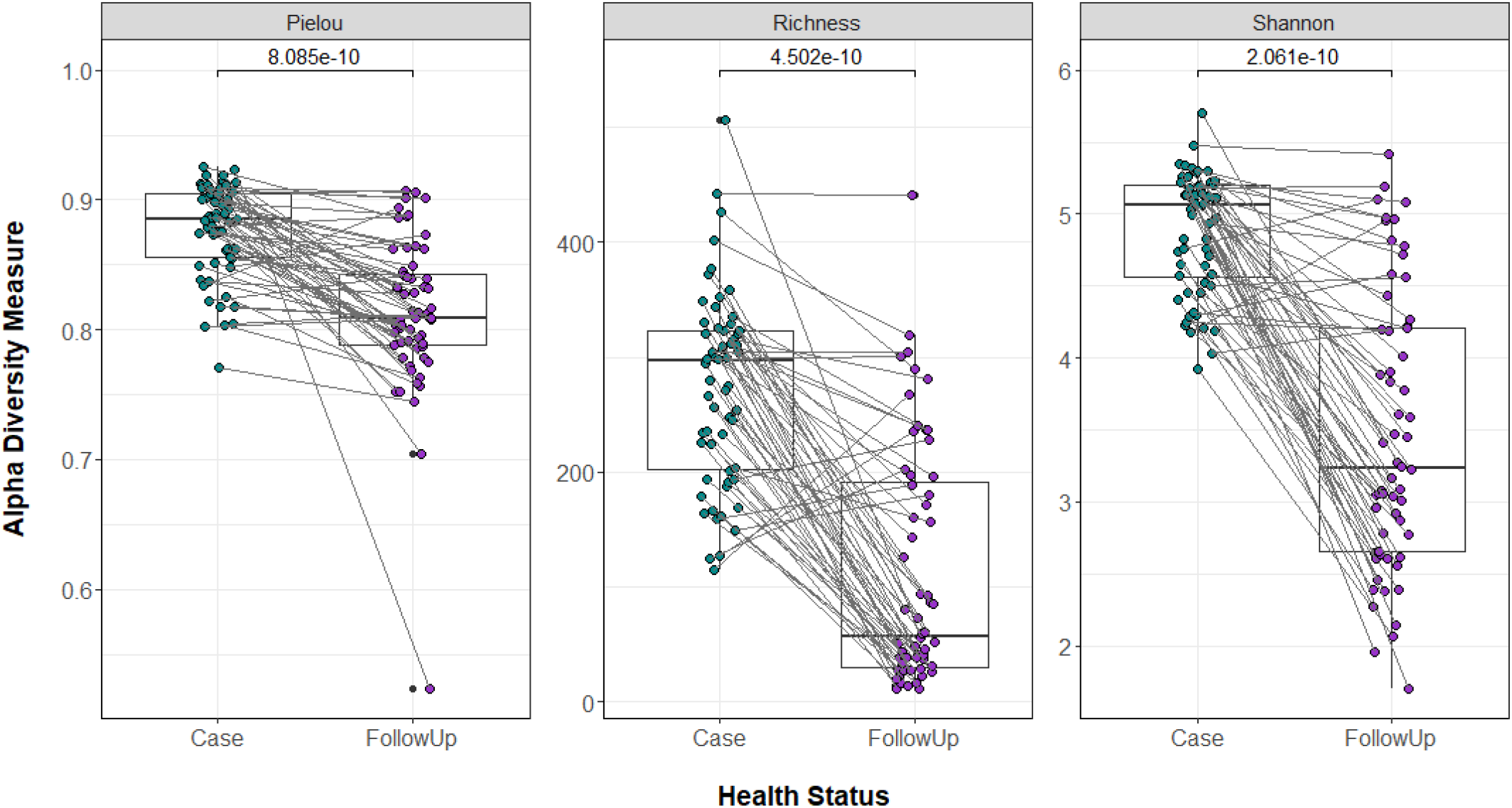
Resistome diversity is greater during infection than after recovery. Three alpha diversity measures (Richness, Shannon’s Diversity Index, and Pielou’s Evenness Index) are presented. Case samples (Case) are indicated with green dots and follow-up samples (FollowUp) are purple. Points are slightly offset from the vertical to allow interpretation of all samples. The median of each measure is indicated by the thick bar within each box and the first and third quartiles are indicated at the bottom and top of the box, respectively. Gray lines between points connect both samples from the same individual. P-values were calculated using the Wilcoxon signed-rank test for paired samples and are shown above the comparison bar within each plot.

The resistome composition also differed during and after infection as was demonstrated in the PCoA based on the Bray-Curtis dissimilarity (PERMANOVA p=0.000999; F=38.75) (**Figure 2**). No difference was observed in the level of dispersion between groups (PERMDISP p=0.52; F=0.468). The samples from those reporting antibiotic use did not cluster separately from those without antibiotics. Data for residence type, antibiotic use, gender, age, hospital, county of origin, stool type, sequencing run, and number of days between samplings were fit to the ordination. Age in years (p=0.013) and year of collection (p=0.043) independently influenced the distribution of points, whereas residence location, hospital, and the number of days since infection only trended toward significance. The pathogen responsible for the acute infections did not have a significant effect on alpha or beta diversity trends (**Additional file 9**).

**Figure 2.**
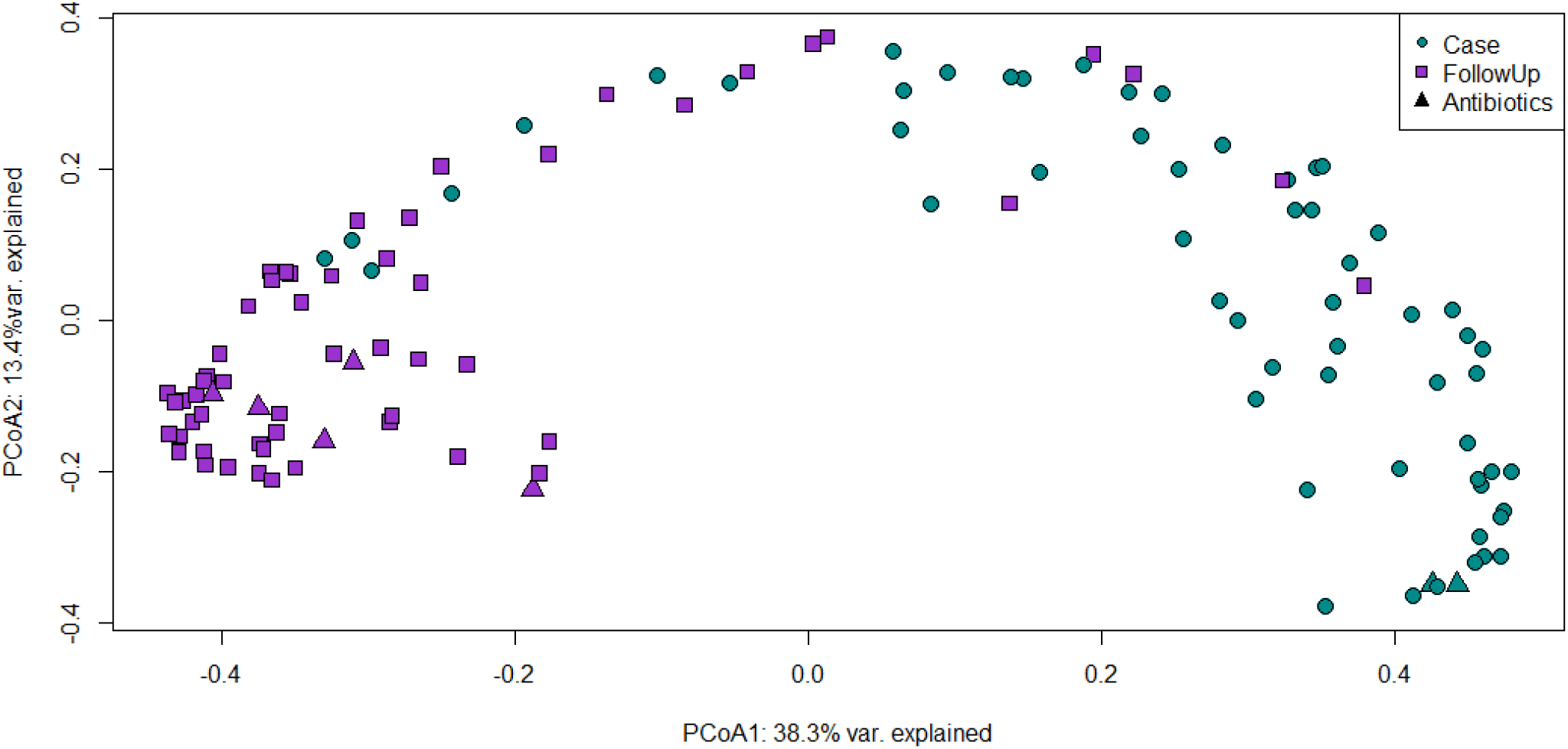
Resistome composition differs significantly during and after infection. A Principal Coordinates Analysis (PCoA) plot of case (green circles) and follow-up (purple, squares) resistomes based on Bray-Curtis dissimilarity calculated from gene-level abundances. The first and second coordinates include the corresponding percentage of similarity explained. Patients that used antibiotics two weeks prior to sample collection are indicated by triangular data points.

### Changes in microbiome composition and diversity post-recovery

A total of 40,022 species, 4,851 genera, 1,157 families, 537 orders, 236 classes, and 224 phyla was found in all samples combined. Notably, the follow-up samples had more diverse gut microbiomes than the cases (**Figure 3**) with significantly greater mean species richness (S_cases_=3,426, S_follow-ups_=5,789; p=2.5e-08), mean evenness (J’_case_=0.150, J’_follow-up_=0.190; p=9.8e-06), and Shannon Diversity (H_cases_=1.21, H_follow-ups_=1.65; p=1.3e-06). When compared to control samples, the follow-ups had similar Shannon Diversity and evenness, though the richness differed (S_follow-ups_=5,789, S_controls_=6,872; p=0.012, Wilcoxon rank-sum test (two-sided, unpaired); **Additional file 8**).

**Figure 3.**
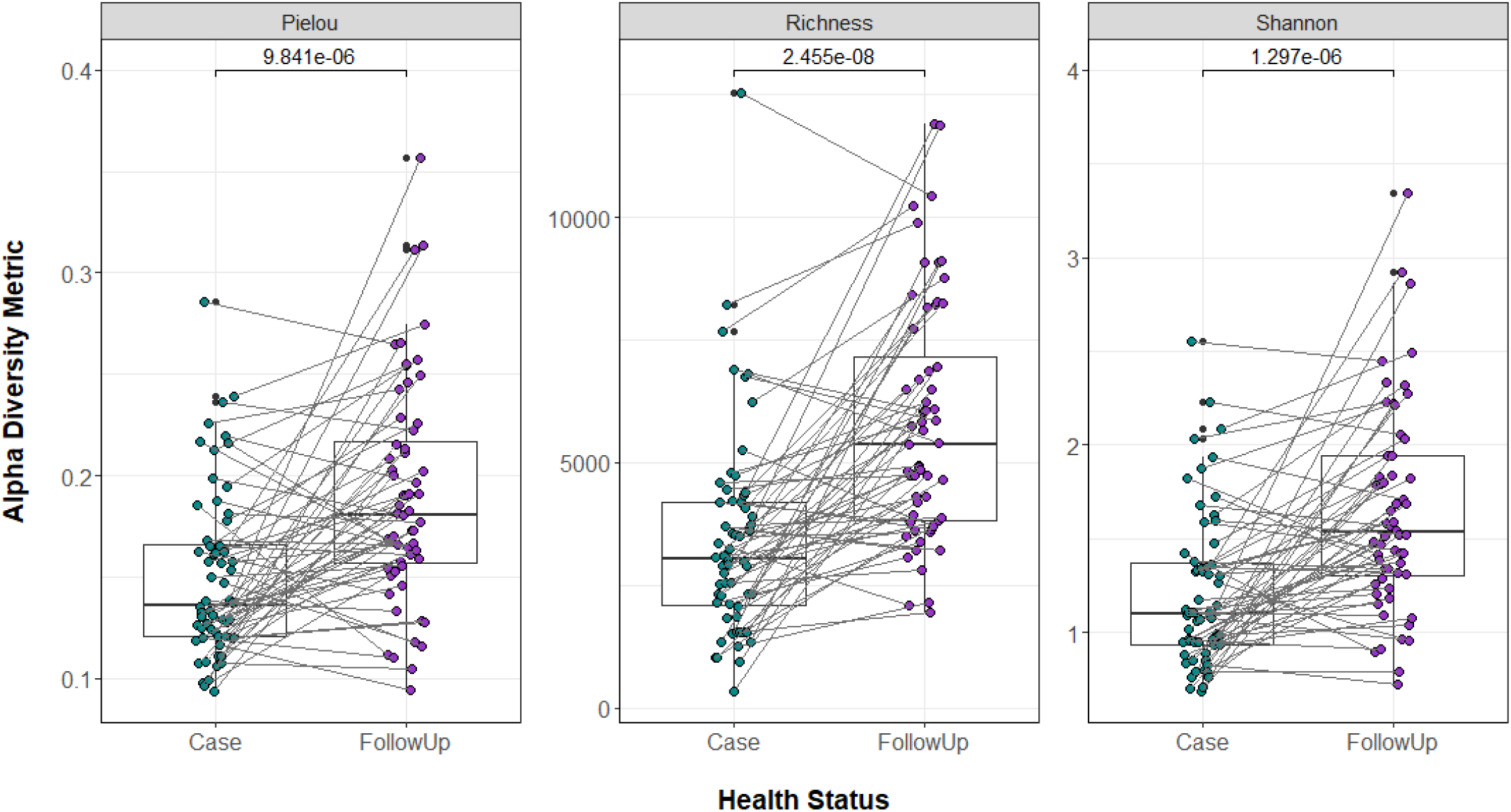
Microbiome diversity is greater after recovery. Box plots show the resistome alpha diversity measures (Pielou’s Evenness Index, Richness, Shannon Diversity Index). Separate points represent case (Case, green) and follow-up (FollowUp, purple) samples and are offset from the vertical for clarity. The median is indicated by the thick bar, while the first and third quartiles are represented by lines at the bottom and top of the box, respectively. Gray lines connect samples from one individual. P-values were calculated using the Wilcoxon signed-rank test for paired samples and are shown above the comparison bars.

The microbiome composition was also significantly different in the case and follow-up samples (PERMANOVA p=0.000999, F=7.31; **Figure 4**), though no difference in the dispersion of points between groups was observed (PERMDISP p=0.086; F=2.86). The same extrinsic covariates were fitted to the PCoA. Age (p=0.008), sequencing run (p=0.001), average genome size (p=0.001), number of genome equivalents (p=0.001), year of sampling (p=0.005), days to follow-up (p=0.013), hospital (p=0.030), and antibiotic use (p=0.008) significantly impacted the point distribution. Similar to the resistome analysis, no differences were observed across pathogens (**Additional file 10**).

**Figure 4.**
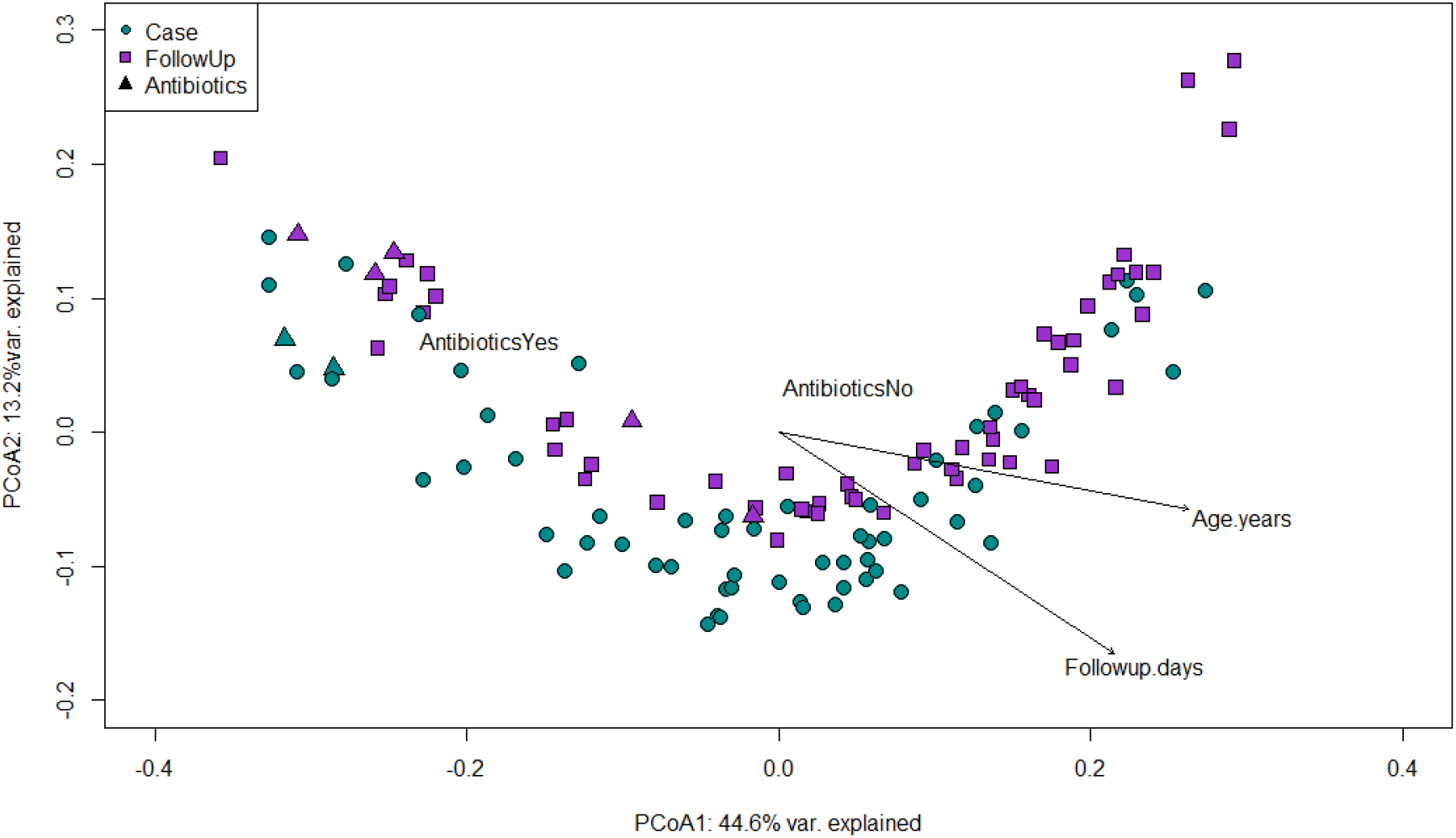
Microbiome compositional differences between cases and follow-ups are nuanced. A Principal Coordinates Analysis plot is shown for case (Case, green circles) and follow-up (FollowUp, purple squares) microbiomes based on Bray-Curtis dissimilarity at the species level. A biplot was overlaid to display variables that had a significant influence on the distribution of points in the ordination. Age (Age.years), number of follow-up days (Followup.days), antibiotic use (Yes and No) were influential vectors. The first and second coordinate are shown and include the corresponding percentage of similarity explained. Patients that self-reported use of antibiotics two weeks prior to sample collection are indicated by triangular data points.

### ARG composition and abundance varied during and after infection

The relative abundance of ARGs differed between groups (**Additional file 11**). The top-three resistance classes in cases accounted for 39.8% of the total resistance genes relative to 71.0% for follow-ups, supporting the observation of greater resistome diversity during infection. Classes for drugs and biocides (15.1%), MLS (13.3%), and multi-metals (11.3%) were most abundant in cases compared to MLS (33.5%), tetracyclines (22.0%), and aminoglycosides (15.5%) in the follow-ups (**Figure 5**). In the differential abundance analysis, classes for multi-metal resistance (coef= −0.243; q-value=1.04e-04), drug and biocide resistance genes (coef= − 0.243; q-value= 1.46e-03), drug, metal, biocide resistance (coef=-0.212; q-value=7.86e-09), and fluoroquinolone resistance genes (coef= −0.168; q-value= 8.19e-10) were more abundant in cases (**Additional file 12**). Comparatively, tetracycline resistance genes (coef=0.352; q-value=2.26e-05) were more abundant in the follow-up samples followed by MLS (coef=0.251; q-value=1.49e-25) and aminoglycoside (coef=0.118; q-value= 7.86e-09) genes.

**Figure 5.**
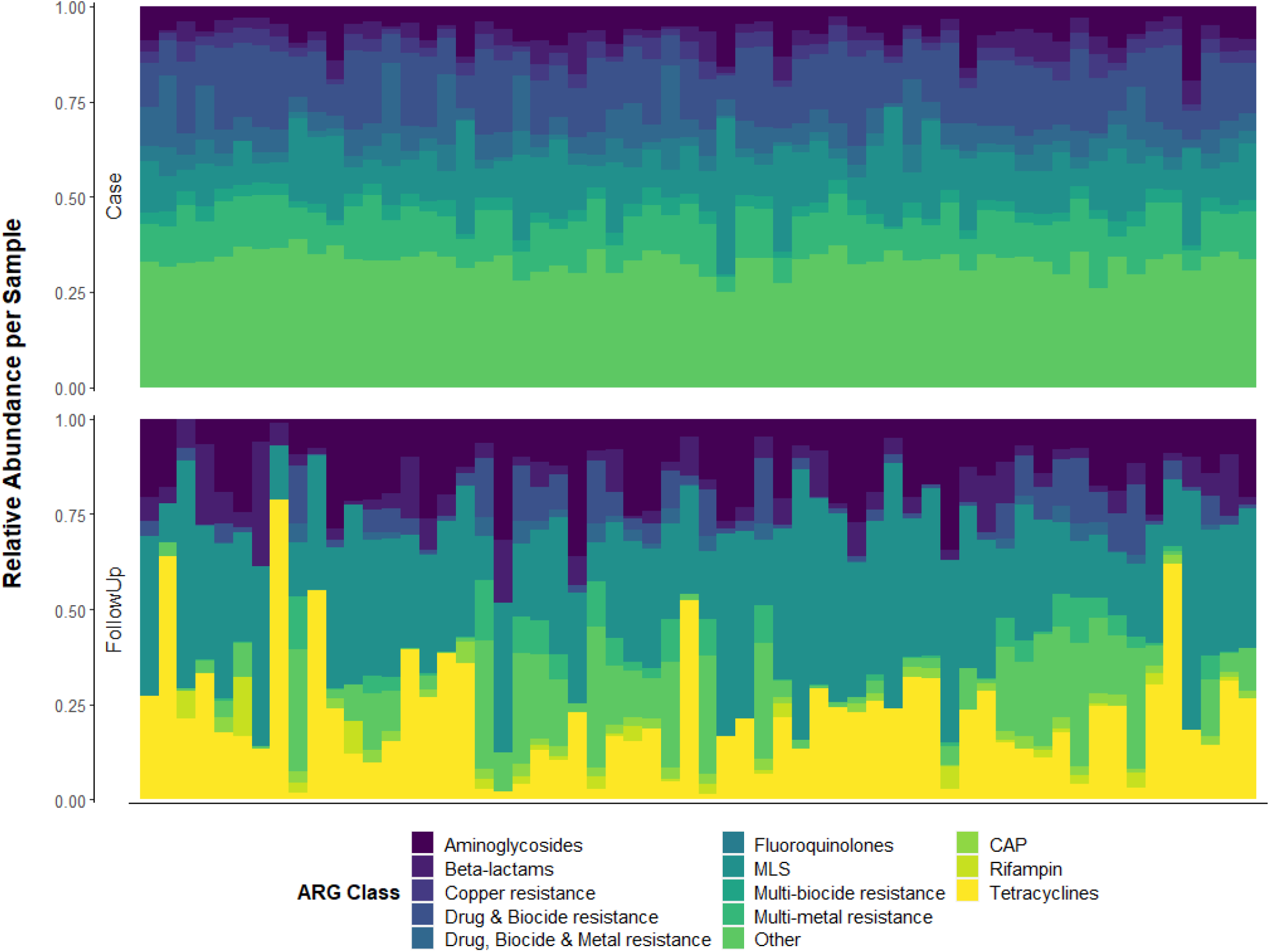
Relative abundance of the top-10 resistance gene classes differs between case and follow-up samples. The top-10 most abundant compound classes is shown for cases (Case, top panel) and follow-ups (FollowUp, bottom panel). Each column represents the resistome from one individual and columns are ordered by the paired samples, meaning that the column position in each side of the plot refers to the same individual during or after infection. Relative abundances were determined using raw gene abundances normalized by the approximate number of genome equivalents in the sample as determined using MicrobeCensus [21]. CAP = cationic antimicrobial peptides; MLS = Macrolide, Lincosamide, Streptogramin; MDR = Multidrug resistance; QACs = Quaternary Ammonium Compounds.

At the group level, specific ARGs were identified for the predominant classes. In the cases, the most abundant groups were MLS23S (11.9%) conferring MLS resistance, *rpoB* (2.8%), a rifampin resistance gene, and A16S (3.8%), which is important for aminoglycoside resistance (**Additional file 13**). Similarly, the differential abundance analysis detected MDR genes, *rpoB* (coef= −0.123; q-value=6.30e-05) and *mdtC* (coef= −0.103; q-value=4.97e-09), to be the most differentiating ARG groups for cases (**Additional file 12**). Genes such as *parC* (coef= −0.102; q-value= 3.90e-11) and *gyrA* (coef= −0.101; q-value=7.38e-08), that encode resistance to fluoroquinolones, were also more abundant in cases.

In the follow-ups, the most abundant groups were for MLS, tetracycline, and aminoglycoside resistance, with MLS23S (n=6.6; 24.3%), *tetQ* (n=4.0; 17.0%), A16S (n=2.4; 9.5%), and *cfx* (n=0.84; 3.8%) predominating, respectively (**Additional file 13**). *tetQ* had the greatest differential abundance in favor of follow-ups (coef=0.30; q-value=6.56e-05) (**Additional file 12**). Despite its noted prevalence among cases, MLS23S was also a defining group for follow-ups since it comprised a greater proportion of ARGs (coef=0.172; q-value=5.54e-06). The *cfx* (coef=0.124; q-value=0.0078) and other genes important for MLS resistance such as *mefE* (coef=0.08; q-value=3.54e-07) and *ermF* (coef=0.07; q-value=3.68e-08), were also more abundant in the follow-ups as were aminoglycoside resistance genes *ant(6*) (coef= 0.103; q-value=5.23e-04) and A16S (coef= 0.092; q-value=5.14e-04).

### Taxa composition and abundance differ markedly during and after infection

Although both cases and follow-ups were dominated by Bacteria (relative abundance = 82.0% and 84.4%, respectively) with fewer Archaea or Eukarya, the members of this kingdom comprising the respective microbiomes were distinct. During infection, cases had a high proportion Proteobacteria (37.1%) with decreased abundance of Bacteroidetes (29.6%) and Firmicutes (13.7%) (**Figure 6**). It is notable that a large proportion of reads could not be assigned to the Phylum level for both the case (16.4%) and follow-up (13.5%) samples. In the differential abundance analysis, Proteobacteria strongly represented cases as well (coef= −0.461; q-value=9.35e-28).

**Figure 6.**
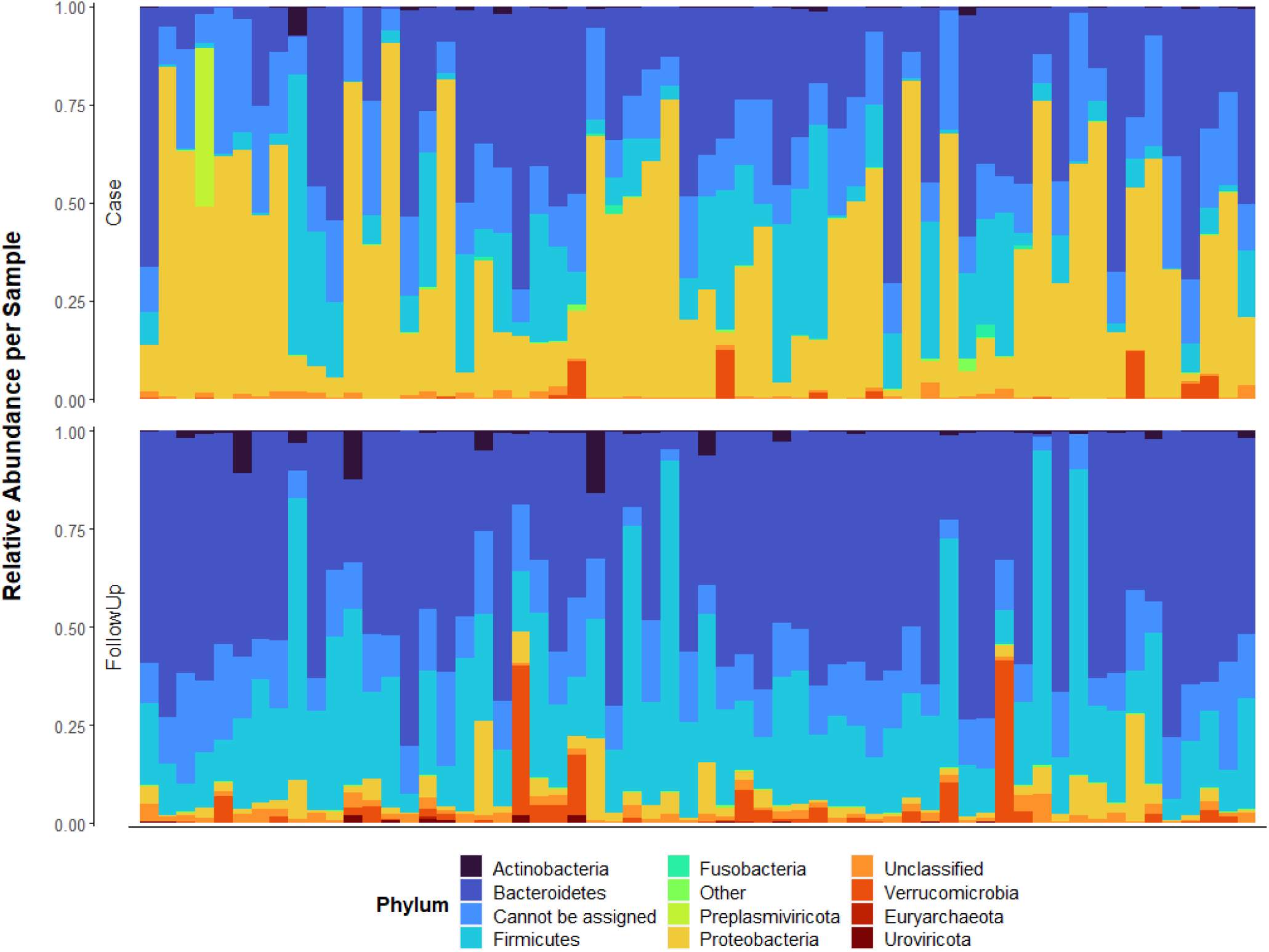
Relative abundance of microbial phyla differs between cases and follow-ups. The top-10 microbial phylum with the greatest average relative abundance among cases (Case, top panel) or follow-ups (FollowUp, bottom panel) is shown; each column represents the microbiome from one individual. Columns are ordered by their sample pairing (i.e., the column position in each plot corresponds to the same individual. Relative abundances were determined using raw gene abundances that had been normalized by the approximate number of genome equivalents in the sample as determined using MicrobeCensus [21].

At the genus level, cases and follow-ups each had high proportions of reads that could not be assigned to a specific genus (case=50.1%; follow-up=46.9%). Beyond this, *Bacteroides* was the most prevalent in both the cases and follow-ups (14.5% and 18.7%, respectively). In cases, this followed by two prominent members of the *Enterobacteriaceae* family within Proteobacteria: *Salmonella* (7.1%) and *Escherichia* (5.0%) (**Additional file 14**). The next highest relatively abundant genus in cases was *Pseudomonas* (2.8%), which is also a member of Proteobacteria. In concordance with these findings, the differential abundance analysis identified *Escherichia* (coef= −0.156; q-value=0.0021) as the predominant genus among the cases, which is mainly represented by *Escherichia coli* (coef=-0.146; q-value=0.0082) (**Additional file 15**). Moreover, *Shigella* (coef= −0.057; q-value=0.0059), which was represented by three species (*S. sonnei, S. flexneri*, and *S. dysenteriae*), as well as *Enterobacter* (coef= −0.020; q-value= 1.10e-08) and *Citrobacter* (coef= −0.017; q-value= 8.07e-06) were also more abundant in the cases.

In follow-ups, both the Bacteroidetes and Firmicutes populations appeared to rebound during recovery and were notably more prevalent (49.3% and 26.9%, respectively). These phyla also defined follow-ups in the differential abundance analysis (Bacteroidetes (coef=0.305; q-value=1.87e-05); Firmicutes (coef=0.199; q-value= 4.61e-07)). Specifically, increases in beneficial genera such as *Alistipes* (5.0%) and *Prevotella* (2.5%) from the Bacteroidetes phylum were observed. The differential abundance analysis, however, detected Firmicutes genera comprising *Roseburia* (coef=0.050; q-value=6.28e-05), *Dialister* (coef=0.038; q-value=0.0036), and *Ruminococcus* (coef=0.037; q-value=2.83e-06) to predominate. *Phocaeicola* was the most abundant genus of Bacteroidetes (coef=0.037; q-value=1.82e-08), which was represented by *Phocaeicola vulgatus* and *Phocaeicola dorei*. One consistent finding among both methods is the heightened abundance of *Akkermansia* (2.8%) from phylum Verrucomicrobia, a defining genus of follow-ups (coef=0.033; q-value= 0.0069).

### The continuous structure of resistome and microbiome compositions

For each level examined (e.g., phylum, genus, ARG class, ARG group), the top contributing features were determined and relevant continuous structure scores were overlaid onto ordination plots using MMUPHin [33]. When considering taxonomy, a tradeoff was observed between the case dominant Proteobacteria phyla and Bacteroides and Firmicutes, which were most abundant in follow-ups and only a subset of cases. At the genus level, an evident gradient was observed between samples containing *Escherichia*, *Salmonella*, *Klebsiella*, *Shigella*, and *Pseudomonas* versus those dominated by *Bacteroides* and *Alistipes* (**Figure 7A**). These differences are visible when overlaid onto ordination as a gradient relevant to loading score (**Figure 7B**). At the species level, which reveals gradients at the greatest resolution, we observed a tradeoff between harboring *Escherichia coli, Klebsiella pneumoniae*, and *Shigella sonnei* versus many *Bacteroides* species including *B. fragilis, B. stercoris, B. uniformis*, etc., and *Phocaeicola* species such as *P. vulgatus* and *P. plebeius* (**Additional file 16**).

**Figure 7.**
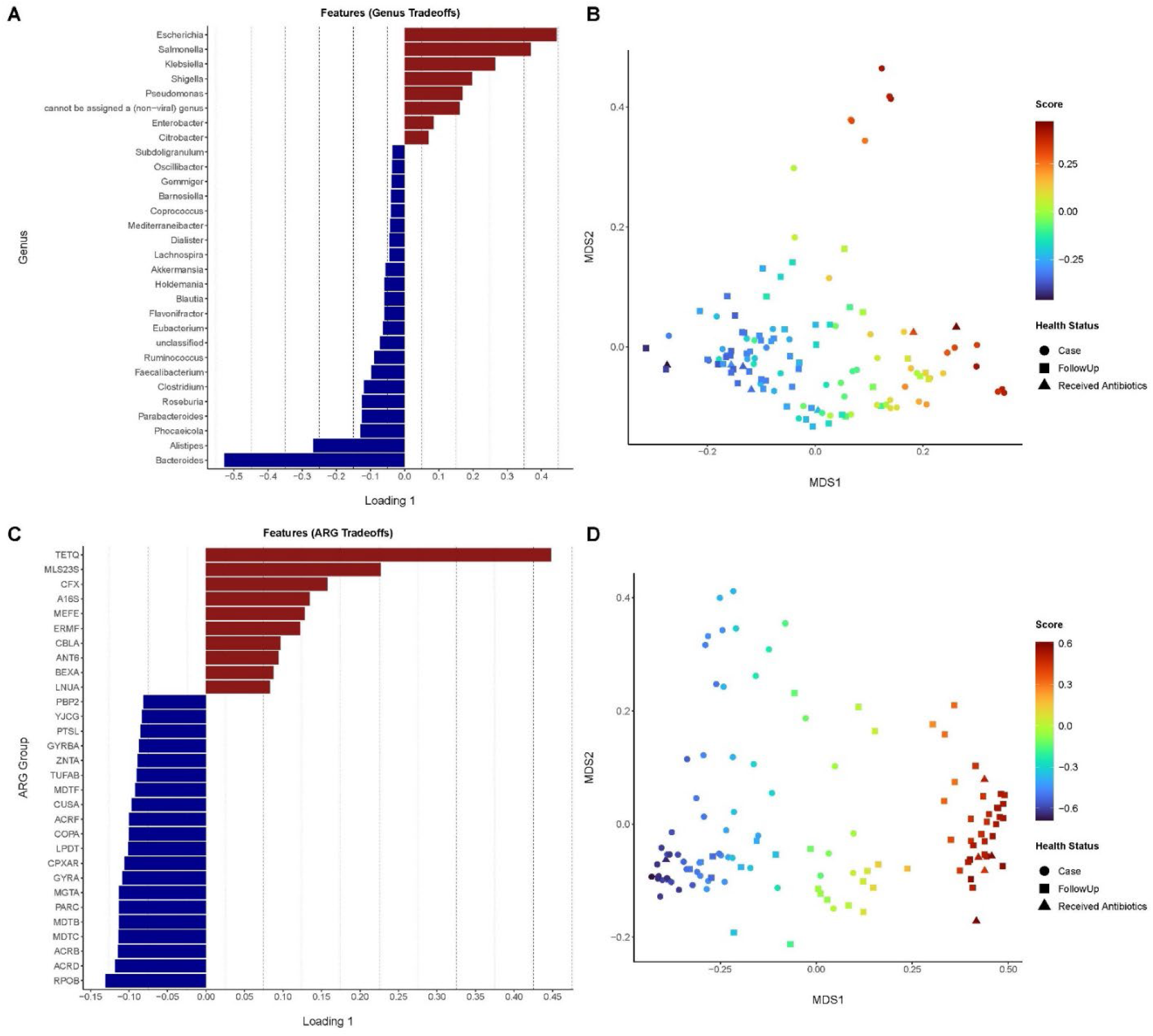
Continuous structure analysis reveals gradients driving the distribution of samples across the population. The top consensus loadings of the PCA for **A)** genus and **C)** antibiotic resistance gene (ARG) groups are shown stratified by sample for cases (Case=red) and follow-ups (FollowUp=blue) drawn from the differential abundance analyses. The composition gradients at the **B)** genus and **D)** ARG group levels overlaid onto ordination plots based on Bray-Curtis dissimilarity. Cases (circles), follow-ups (squares), and individuals who received antibiotics (triangles) are shown. The color gradient (“Score”) refers to the continuous structure score affiliated with Loading 1 for phyla and genera, respectively. Juxtaposition of (A-C) and (B-D) allow interpretation of tradeoffs within the samples.

Tradeoffs were also observed for different resistance genes. At the class level, there was a continuous gradient relative to tetracycline, MLS, and aminoglycoside dominant resistomes versus ARGs for multi-metal resistance, drug and biocide resistance, and drug, metal, and biocide resistance classes (data not shown). At the ARG group level, *tetQ* was identified as a dominant driver of continuous structure scoring for follow-ups (**Figure 7C**), whereas resistance genes such as *rpoB, acrA, acrB, mdtC*, and *mdtB* were defining for the opposite side of the PCoA axis. Overlaying these loading scores onto ordination further revealed the taxonomic gradients among case and follow-up samples (**Figure 7D**).

### Different ARG-harboring microbial hosts are present in cases and follow-ups

In cases, ACCs, on average, were primarily attributed to *Escherichia* (38.05%) followed by *Salmonella* (18.31%) and *Klebsiella* (9.92%) (**Figure 8**). Of the *Escherichia*-associated ARGs, 27.4% were assigned to MDR on average, though ARGs relevant to drug and biocide resistance (8.12%), fluoroquinolone resistance (7.06%), and aminoglycoside resistance (6.21%) were also identified. Comparatively, the *Salmonella*-associated ACCs mostly contained genes for MDR and drug and biocide resistance (16.5% and 11.7%, respectively), while the *Klebsiella* ACCs harbored an array of fosfomycin resistance genes (13.3%) followed by transposase genes in the IS5 family (12.6%). *Klebsiella* ACCs also contained ARGs for elfamycin resistance (10.4%) and MDR (9.08%).

**Figure 8.**
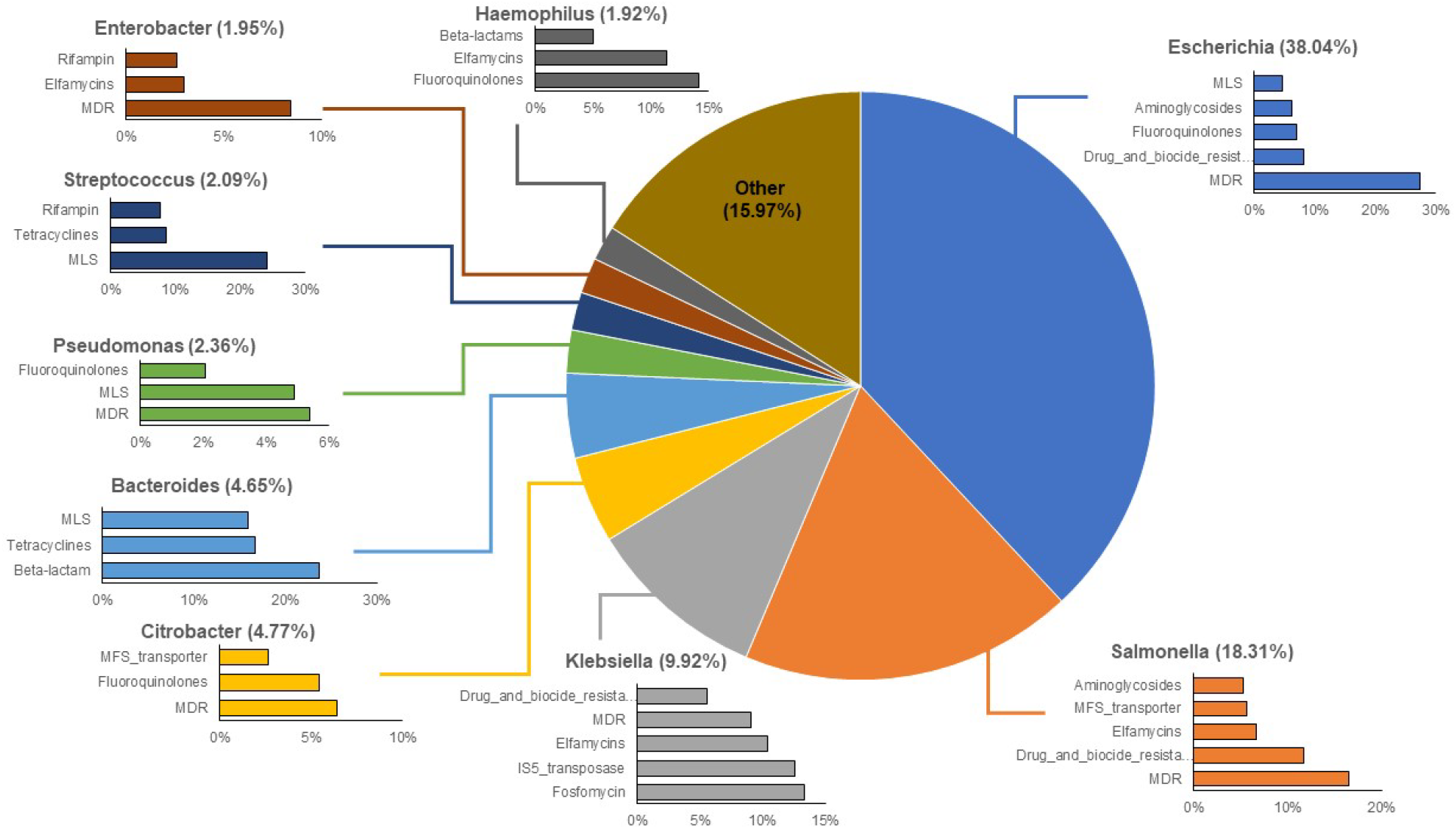
The top-10 genera assigned to antibiotic resistance gene (ARG)-carrying contigs (ACCs) in case samples. The percentages associated with each genus indicate the percent of ACCs assigned to that genus. Each bar chart associated with a genus displays the top-5 or top-3 ARG classes affiliated with that particular genus on the ACCs.

Although the most prominent genus in follow-up ACCs was also *Escherichia* (19.81%), the next most prevalent genera were classified as *Bacteroides* (15.12%) and *Faecalibacterium* (5.99%) (**Figure 9**). Notably, the array of ARGs harbored in the *Escherichia*-associated ACCs was nearly identical to cases with MDR genes predominating (25.1%), followed by resistance to drugs and biocides (4.71%), fluoroquinolones (4.70%), and aminoglycosides (3.84%). Of the *Bacteroidetes-associated* ACCs, genes for MLS, beta-lactam, and tetracycline resistance were the most common. The 5.21% of the ACCs that could not be classified and represented an “Uncultured” taxon harbored ARGs for tetracyclines, beta-lactams and phenicols.

**Figure 9.**
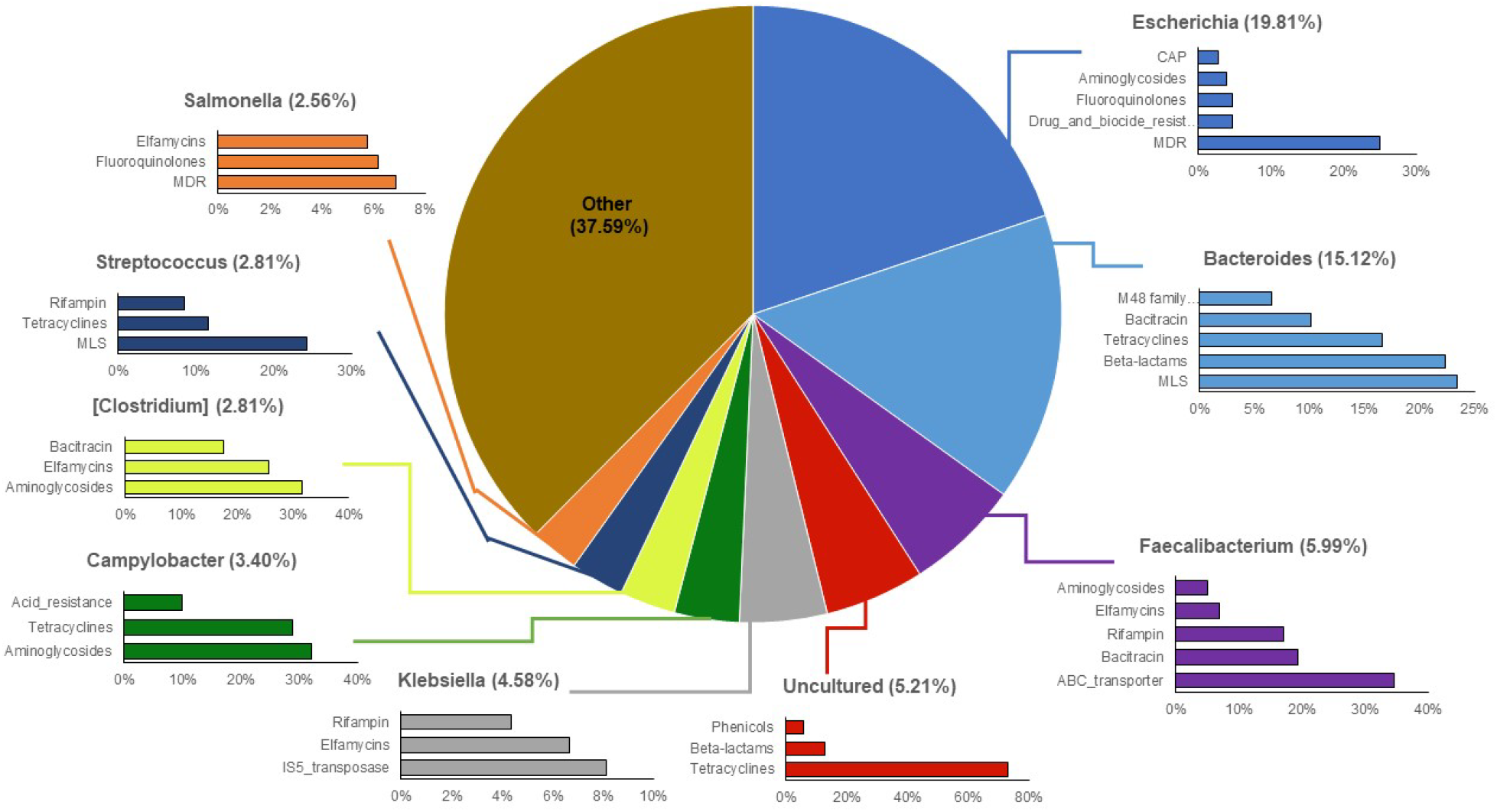
The top-10 genera assigned to antibiotic resistance gene (ARG)-carrying contigs (ACCs) in follow-up samples. The percentages associated with each genus indicate the percent of ACCs assigned to that genus. Each bar chart associated with a genus displays the top-5 or top-3 ARG classes affiliated with that particular genus on the ACCs.

### Microbes linked to case infections harbor ARGs during and after recovery

Differences in ACCs were also identified after stratifying by the bacterium linked to each infection. Among the 23 cases with *Campylobacter* (n=23) infections, for instance, the genera comprising the greatest proportion of ACCs were *Escherichia* (42.84%), *Klebsiella* (10.01%), and *Salmonella* (7.09%). Upon recovery, however, *Campylobacter* cases most often had ACCs representing *Bacteroides* (18.34%), followed by *Escherichia* (17.31%) and *Faecalibacterium* (6.76%). It is also notable that *Campylobacter* was in the top-20 genera represented on ACCs, with proportions of 1.96% and 3.81% in cases and follow-ups, respectively. ARGs harbored by *Campylobacter* in case samples conferred resistance to tetracyclines (27.6%), aminoglycosides (9.92%), and rifampin (8.31%). Among the *Campylobacter* ACCs in follow-up samples, tetracycline (29.0%) and aminoglycoside (27.0%) resistance genes were commonly detected as were genes for MLS resistance (10.3%). Importantly, genes encoding resistance to aminoglycosides were 2.7 times more prevalent among *Campylobacter* ACCs in the follow-up samples relative to the case samples.

In the 29 cases with *Salmonella* infections, most ACCs were classified as *Escherichia* (32.39%)*, Salmonella* (30.96%), and *Klebsiella* (7.89%) compared to *Escherichia* (20.66%), *Bacteroides* (14.16%), and *Faecalibacterium* (6.29%) for the follow-up samples. The most common genes detected in *Salmonella* ACCs were important for multi-compound resistance including drug and biocide resistance (14.1%) and MFS transporters (13.1%), which can have MDR effects or high specificity to certain classes. ARGs for drug, biocide, and metal resistance (7.61%) were also identified. Among the follow-up samples, the most prevalent class within *Salmonella*-associated ACCs was RND efflux transporters (9.29%), followed by MFS transporters (6.84%) and fluoroquinolone resistance genes (6.31%).

### Multiple clinically relevant ESBLs can be found after recovery from enteric infections

In all, 49 distinct genes encoding beta-lactam resistance were identified representing class A, C, and D beta-lactamases (**Additional file 17**); 11 (22.4%) were classified as distinct genes encoding ESBL production that confer resistance to multiple beta-lactam antibiotics. Among the ESBL genes detected, those belonging to the CepA family of class A beta-lactamases were most prevalent occurring in 19 case and 13 follow-up samples; each gene was taxonomically assigned to *Bacteroides*. Of these 19 cases, the gene was absent or “lost” in nine patients at follow-up despite being detected in the case sample (**Additional file 18**). For the remaining 10 cases, the same gene was detected in both the case and follow-up samples, indicating persistence with the *Bacteroides* population. Additionally, three patients acquired a gene during the recovery period as it was present in follow-up sample but not the initial sample collected during infection.

ESBL genes of the OXA family, which included OXA-1, OXA-50, OXA-51, and OXA-61, were also detected; however, each gene was attributed to a different microbial host in the ACC analysis and was only found in 2-3 individuals. Although the OXA-61 family of class D beta-lactamases was harbored by *Campylobacter*, it was only found in two of the 23 cases with *Campylobacter* infections. Similarly, *Klebsiella* was found to harbor OXY genes in four case samples, though these were absent in the follow-up samples. *Klebsiella* also possessed genes representing the SHV family of class A beta-lactamases, which were detected in eight cases. Because SHV genes were also detected in two unpaired follow-ups, it is likely that all eight cases lost the genes and two patients acquired it during recovery.

Genes representing the ADC family of class C ESBLs harbored by *Acinetobacter* were also detected, though there was not enough evidence to infer transfer of these genes between taxa. Since many of the ESBLs were present in cases but not follow-ups, we could not assess whether they were transferred horizontally among bacteria during recovery. Other relevant beta-lactamases were also identified including the BlaEC family of class C beta-lactamases, which were primarily attributed to genus *Escherichia* and were found in 49 cases and 19 follow-ups. Intriguingly, the ARG was lost in 35 cases, maintained in 14, and acquired in 5 follow-ups.

Genes encoding the CfxA family of class A broad-spectrum beta-lactamases were also detected and were primarily harbored by *Bacteroides*, but also appeared within *Prevotella*. Among these *Bacteroides*-associated ARGs, 46 were found in cases and 48 in follow-ups. Although only 7 of these genes were lost by cases, 39 were maintained and 9 were acquired during recovery. A similar trend was observed for *cfxA* within *Prevotella* as three of the five cases lost the genes, two maintained them, and seven acquired them during recovery. Interestingly, there is evidence of horizontal transfer of these CfxA genes between *Bacteroides* and *Prevotella*. For example, six separate case-follow-up pairs show *cfxA* as being “acquired” by *Prevotella* in follow-ups but also maintained by *Bacteroides*, suggesting potential *Bacteroides*-to-*Prevotella* transfer. Two other case-follow-up pairs had *cfxA* maintained in both *Bacteroides* and *Prevotella* during recovery, while there were three instances in which the *Prevotella*-harbored ARG was “lost” and the *Bacteroides*-harbored *cfxA* was maintained, suggesting the possibility of *Prevotella-to-Bacteroides* transfer.

Genes encoding the broad CMY-family of class C beta-lactamases were also identified and assigned to *Salmonella* in 3 cases (all lost) and 2 follow-ups (both acquired). Relatedly, the CMY-2 family of class C beta-lactamases was identified within *Citrobacter* and *Salmonella*. Among these ARGs harbored by *Citrobacter*, 8 were found in case samples and 3 in follow-ups; 6 cases lost the gene, 2 maintained it, and one follow-up acquired it. Of the CMY-2 ARGs harbored by *Salmonella*, two were found in cases (each of which were lost) and one was acquired in a follow-up sample. Although the CMY family is a broader category than the CMY-2 family of beta-lactamases, it is possible that the CMY family defined in our study contains CMY-2 genes relevant to this analysis. For example, there is one case-follow-up pair in which the CMY-2 family was maintained in *Citrobacter* and the CMY family was acquired in *Salmonella;* yet another case-follow-up pair indicated loss of the CMY family of beta-lactamases in *Salmonella* but maintenance and noted increase of the CMY-2 family in *Citrobacter*. Although loosely inferred, these data indicate the potential for the horizontal transfer of CMY-family genes across genera.

Finally, genes for the general subclass A2 of class A beta-lactamases were found in *Bacteroides* among both the cases (n=45) and follow-ups (n=47); 7 cases lost the gene during recovery, while 38 maintained it and 9 follow-ups acquired it. The more general “class A beta-lactamase” gene was also found in nine other genera including *Atlantibacter*, *Bacillus, Burkholderia, Clostridium, Proteus, Salmonella, Yersinia, Escherichia*, and *Klebsiella*. Although there is a slight difference in resolution of these identified features, it is helpful to consider the potential for transfere across genera.

## DISCUSSION

The human gut microbiome, when disrupted by an infectious pathogen, can drastically change in composition taxonomically, genetically, and functionally [35]. In most instances, pathogen invasion leads to a state of dysbiosis linked to a decrease in gut microbiota diversity [4, 36]. Our study supports these findings, as markedly lower microbiome diversity was observed among cases during infection than after recovery regardless of the bacterial pathogen causing infection. The observed shifts in microbiome composition post-recovery are indicative of gut health, as healthy family members (controls) and follow-ups had more similar microbiome profiles than the cases. In addition to the increased microbiota diversity post-recovery, specific taxonomic signatures such as enhanced abundance of Bacteroidetes and Firmicutes, were observed. For instance, members of *Bacteroides, Prevotella*, and *Phocaeicola* as well as *Faecalibacterium, Roseburia*, and *Ruminococcus* were found, which have been shown to play influential roles in maintaining gut homeostasis and metabolic health [37–39]. By contrast, the cases were defined primarily by members of Proteobacteria such as *Escherichia, Salmonella, Shigella*, and *Klebsiella*, which have been linked to acute enteric disturbances as well as prolonged dysbiosis and long-term disease outcomes [40].

The opposite was true for the collection of ARGs, as cases had greater resistome diversity during infection than after recovery. Because shifts in microbial composition inherently influence the presence and abundance of ARGs harbored by microbes within a community, this finding is not surprising. Among the key differences observed, cases had more multi-compound and multi-drug resistance genes during infection than post-recovery, whereas tetracycline, MLS, and aminoglycoside resistance genes were more abundant in the recovered (follow-up) sample. Diverse sets of ARGs have previously been found in otherwise healthy individuals as well [10, 41, 42], providing additional support for the human gut as an important reservoir of antibiotic resistance determinants [14].

Intriguingly, a subset of five follow-up samples were more closely related to the case microbiome and resistome samples in the PCoA. Because these patients had an average number of 110 days since infection, which did not differ from the overall mean (n=108 days), other factors likely contributed to the case-like microbiome profiles observed. Indeed, four patients were either <10 or >50 years of age and two of these individuals were hospitalized. Since children and older individuals typically have an enhanced risk of developing more severe disease [43, 44], these patients could have experienced lengthier infections than other members of the sample cohort. The same is true for those who were hospitalized and hence, the microbiome may have not fully recovered at the time of follow-up sampling. The complete level of microbiome recovery, however, could not be deduced for any of the patients since we did not evaluate the gut microbiome in the same patients prior to infection. It is likely that the state of the microbiome prior to infection as well as its resilience to disturbances will vary across individuals and greatly impact the trajectory of disease and recovery. Implementation of a more rigorous longitudinal study is therefore needed in the future.

In the host-tracking analysis, we demonstrated that specific microbial taxa were more likely to harbor ARGs during infection. *Escherichia*, for instance, was a prominent host in the cases regardless of the pathogen linked to the infection. Specifically, *Escherichia* comprised an average of 38% of all ACCs, with most genes being important for MDR or multi-compound resistance. This result is not surprising given the increased abundance of *Escherichia* observed during infection. Expansion of *Escherichia* and Enterobacteriaceae in general, was previously suggested to be linked to inflammation in the gut [45], which was also shown to augment HGT rates between commensal and pathogenic members of this family [46]. Moreover, as the level of MDR increases within a population, so too does the number of integrons, which were also shown to persist among commensal *E. coli* [47]. This enhanced mobility and maintenance of resistance determinants are key contributors to the emergence of resistant pathobionts [3, 48].

Evidence of ARGs harbored by genera linked to the acute infections was also observed, indicating that some pathogens bring resistance genes into the gut during infection. In patients with *Salmonella* infections, for instance, *Salmonella* accounted for ~31% of all ACCs compared to the overall case average of 18%, with most genes encoding MDR or drug and biocide resistance. Co-selection for resistance to antibiotics, metals, and biocides has been previously documented in *Salmonella* and other foodborne pathogens [49]. This evidence is supported by data generated in a co-occurrence network analysis despite being a less robust approach [50]. Notably, a *Salmonella*-specific subnetwork comprised of multiple metal, biocide, and MDR genes was identified among *Salmonella* cases (**Additional file 20**). This subnetwork was not detected in the co-occurrence network generated using data from the *Campylobacter* cases alone (**Additional file 21**). Hence, these findings indicate that the different *Salmonella* pathogens brought similar ARGs into the microbial communities at the time of infection. Future whole-genome sequencing studies, however, should be conducted to characterize each pathogen and determine the diversity and frequency of those ARGs that were introduced into each gut community.

In the follow-up samples, *Escherichia* still accounted for the greatest proportion (~20%) of all ARG-carrying contigs, which mostly contained MDR genes; however, the proportion was 1.9 times less than that observed during infection. Unlike the cases, *Bacteroides* was the second most important genus accounting for ~15% of the ARG-carrying contigs at recovery with MLS, beta-lactam, and tetracycline resistance genes predominating. Members of Bacteroidetes and Firmicutes have previously been linked to high levels of tetracycline and erythromycin resistance carrying genes such as *tetQ* as well as *ermF* and *ermG*, respectively [51]. These genes were previously suggested to be maintained in microbial host populations even in the absence of antibiotic selection, thereby enhancing the likelihood of HGT [51]. Although resistance to bet-alactam antibiotics has been documented, variation in resistance rates has been observed across species and geographic locations, particularly for the beta-lactamase producers [52, 53].

Indeed, the transfer and acquisition of genes encoding beta-lactamase production is of great concern. During enteric infection, we detected 11 distinct ESBLs that varied in frequency among the cases, although this number may underestimate the actual diversity as not all sequences could be assigned a class designation. Except for the CepA family of genes, most genes were “lost” or undetectable during recovery. This result is consistent with a prior study showing that some ESBLs including CTX and SHV, were more readily lost, though this was dependent on the bacterial host [54]. The noted roles of *Klebsiella* and *Escherichia* in harboring ESBLs in both the case and follow-up samples calls attention to the documented capacity of these genera to transfer genes across species or clonal lineages [55]. *Klebsiella*, for instance, was a prominent ARG carrier in 9.2% and 4.6% of ACCs in the cases and follow-ups, respectively, and was associated with a high occurrence of the IS5 family of transposases. The identification of a genomic element with the potential to transfer ARGs within the gut microbiome is notable, particularly to other members of *Enterobacteriaceae*, which have contributed to the widespread distribution and spread of ESBL genes [2]. Several beta-lactamase genes were also detected that were not classified as extended spectrum. The CfxA gene family, for example, was harbored by both *Bacteroides* and *Prevotella*. In several paired case/follow-up samples, there is evidence for the transfer of *cfxA* between genera, which has been documented previously [56]. Because this evidence is solely based on the detection of the gene in both genera at two different time points, more rigorous methods, such as characterizing the sequence-level similarity, are required for confirmation.

There are other limitations related to the ACC analysis as well. One example is the potential for misclassifying ARGs found on plasmids even though they were previously shown to contain taxonomic information regarding the host microbe [57]. Because assembly of short-read sequences can inaccurately characterize plasmids and other MGEs [58], deeper sequencing is needed to generate more complete assemblies and avoid misclassifying the microbial hosts. In addition, multiple ARGs were attributed to “uncultured” microbes, highlighting the need for more comprehensive databases that can accurately predict host taxonomies. Finally, the ACC analysis relies on classifying microbial hosts based on the co-occurrence of an ARG and its taxa on the same contig. Alternative methods such as Single-molecule Real-time sequencing, are therefore required in future studies. Despite these limitations, this study provides important data about the most common alterations in the gut microbiome and resistome among patients with enteric infections. It also illustrates how infected microbial communities recover, which is needed to guide the development of more targeted intervention strategies or therapeutic options aimed at restoring the dysbiotic gut. Future work should focus on understanding the trajectory of recovery as it pertains to the presence and dissemination of drug resistance and characterizing the interactions between microbial hosts, ARGs, and MGEs during recovery.

## Declarations

### Ethics approval and consent to participate

Study protocols and consent procedures were approved by the Institutional Review Boards at Michigan State University (MSU; IRB #10-736SM) and the MDHHS (842-PHALAB) as well as the four participating hospital laboratories.

### Availability of data and materials

Sequencing reads were deposited in the National Center for Biotechnology Information (NCBI) sequence read archive (SRA) database under BioProjects PRJNA862908 and PRJNA660443 (BioSamples SAMN29999523 to SAMN29999673 and SAMN15958881 to SAMN15958950, respectively). Bioinformatic scripts are at: github.com/ZoeHansen/PAPER_Hansen_Microbiome_2022.

### Authors’ contributions

SDM and JTR conceptualized the study, obtained funds for the project, and organized sample collection and processing. SDM supervised the study, and ZAH completed the data analyses, figure generation, and developed the first manuscript draft. ZAH, SDM, KV, KTS, and LZ assisted with additional analyses and manuscript revisions. All authors read and approved the final manuscript.

## Acknowledgements

We thank Ben Hutton and Jason Wholehan at the MDHHS for help with specimen processing and culture as well as Rebekah E. Sloup, Katherine Jernigan, and Samantha Carbonell, for help with community DNA isolation and sequencing. We also thank Dr. James M. Tiedje for his ongoing and helpful discussions about these data.

## REFERENCES

1. Scallan E, Hoekstra RM, Angulo FJ, Tauxe RV, Widdowson M-A, Roy SL, et al. Foodborne illness acquired in the United States -- Major pathogens. Emerg Infect Dis. 2011;17(1):7–15.

2. Centers for Disease Control and Prevetion (CDC): Antibiotic Resistance Threats in the United States, 2019. Atlanta, GA; 2019. Available at: https://www.cdc.gov/drugresistance/pdf/threats-report/2019-ar-threats-report-508.pdf

3. Wallace MJ, Fishbein SRS, Dantas G. Antimicrobial resistance in enteric bacteria: current state and next-generation solutions. Gut Microbes. 2020;12(1):e1799654; doi: 10.1080/19490976.2020.1799654.

4. Singh P, Teal TK, Marsh TL, Tiedje JM, Mosci R, Jernigan K, et al. Intestinal microbial communities associated with acute enteric infections and disease recovery. Microbiome. 2015;3:45.

5. Le Chatelier E, Nielsen T, Qin J, Prifti E, Hildebrand F, Falony G, et al. Richness of human gut microbiome correlates with metabolic markers. Nature. 2013;500:541–6.

6. Huang AD, Luo C, Pena-Gonzalez A, Weigand MR, Tarr CL, Konstantinidis KT. Metagenomics of two severe foodborne outbreaks provides diagnostic signatures and signs of coinfection not attainable by traditional methods. Appl Environ Microbiol. 2017;83:e02577–16.

7. Argüello H, Estellé J, Zaldívar-López S, Jiménez-Marín Á, Carvajal A, López-Bascón MA, et al. Early *Salmonella* Typhimurium infection in pigs disrupts microbiome composition and functionality principally at the ileum mucosa. Sci Rep. 2018;8.

8. Haag L-M, Fischer A, Otto B, Plickert R, Kühl AA, Göbel UB, et al. Intestinal microbiota shifts towards elevated commensal *Escherichia coli* loads abrogate colonization resistance against *Campylobacter jejuni* in mice. PLoS One. 2012;7:e35988.

9. Yang J, Chen W, Xia P, Zhang W. Dynamic comparison of gut microbiota of mice infected with *Shigella flexneri* via two different infective routes. Exp Ther Med. 2020; 19: 2273–2281.

10. Hansen ZA, Cha W, Nohomovich B, Newton DW, Lephart P, Salimnia H, et al. Comparing gut resistome composition among patients with acute *Campylobacter* infections and healthy family members. Sci Rep. 2021;11: 22368.

11. Lozupone C. Diversity, stability and resilience of the human gut microbiota. Nature. 2012;489:220–30.

12. Reid G, Howard J, Siang Gan B. Can bacterial interference prevent infection? Trends Microbiol. 2001;9:424–8.

13. Sassone-Corsi M, Raffatellu M. No Vacancy: How beneficial microbes cooperate with immunity to provide colonization resistance to pathogens. J Immunol. 2015;194:4081–7.

14. Salyers AA, Gupta A, Wang Y. Human intestinal bacteria as reservoirs for antibiotic resistance genes. Trends Microbiol. 2004;12:412–6.

15. Ma L, Li B, Jiang X-T, Wang Y-L, Xia Y, Li A-D, et al. Catalogue of antibiotic resistome and host-tracking in drinking water deciphered by a large scale survey. Microbiome. 2017;5:154.

16. Doster E, Lakin SM, Dean CJ, Wolfe C, Young JG, Boucher C, et al. MEGARes 2.0: A database for classification of antimicrobial drug, biocide and metal resistance determinants in metagenomic sequence data. Nucleic Acids Res. 2020;48(D1):D561–D9.

17. Li H, Durbin R. Fast and accurate short read alignment with Burrows-Wheeler transform. Bioinformatics. 2009;25:1754–60.

18. Li H, Handsaker B, Wysoker A, Fennell T, Ruan J, Homer N, et al. The sequence alignment/map format and SAMtools. Bioinformatics. 2009;25:2078–9.

19. Quinlan AR, Hall IM. BEDTools: a flexible suite of utilities for comparing genomic features. Bioinform App Note. 2010;26:841–2.

20. McArthur AG, Waglechner N, Nizam F, Yan A, Azad MA, Baylay AJ, et al. The comprehensive antibiotic resistance database. Antimicrob Agents Chemother. 2013;57:3348–57.

21. Nayfach S, Pollard KS. Average genome size estimation improves comparative metagenomics and sheds light on the functional ecology of the human microbiome. Genome Biol. 2015;16:51.

22. Rodriguez-R LM, Konstantinidis KT. Nonpareil: a redundancy-based approach to assess the level of coverage in metagenomic datasets. Bioinformatics. 2014;30:629–35.

23. Bushnell B. BBMAP. Available at: sourceforge.net/projects/bbmap/.

24. Li D, Liu C-M, Luo R, Sadakane K, Lam T-W. MEGAHIT: an ultra-fast single-node solution for large and complex metagenomics assembly via succinct de Bruijn graph. Bioinformatics. 2015;31:1674–6.

25. Mikheenko A, Saveliev V, Gurevich A. MetaQUAST: Evaluation of metagenome assemblies. Bioinformatics. 2016;32.

26. Eren AM, Kiefl E, Shaiber A, Veseli I, Miller SE, Schechter MS, et al. Community-led, integrated, reproducible multi-omics with anvi’o. Nat Microbiol. 2021;6:3–6.

27. Hyatt D, Chen G-L, Locascio PF, Land ML, Larimer FW, Hauser LJ. Prodigal: prokaryotic gene recognition and translation initiation site identification. BMC Bioinform. 2010:11:119.

28. Menzel P, Ng KL, Krogh A. Fast and sensitive taxonomic classification for metagenomics with Kaiju. Nat Commun. 2016;7:11257.

29. Li Y, Xu Z, Han W, Cao H, Umarov R, Yan A, et al. HMD-ARG: hierarchical multi-task deep learning for annotating antibiotic resistance genes. Microbiome. 2021;9.

30. Buchfink B, Xie C, Huson DH. Fast and sensitive protein alignment using DIAMOND. Nat Methods. 2015;12:59–60.

31. Ma L, Xia Y, Li B, Yang Y, Li LG, Tiedje JM, et al. Metagenomic assembly reveals hosts of antibiotic resistance genes and the shared resistome in pig, chicken, and human feces. Environ Sci Technol. 2016;50:420–7.

32. Oksanen J, Blanchet FG, Friendly M, Kindt R, Legendre P, McGlinn D, et al. Package ‘vegan’: Community Ecology Package: Ordination, Diversity and Dissimilarities 2019;2(9). Available at: https://cran.r-project.org/web/packages/vegan/vegan.pdf

33. Ma S: MMUPHin: Meta-analysis methods with uniform pipeline for heterogeneity in microbiome studies. Available at: https://rdrr.io/bioc/MMUPHin/

34. Lin H, Peddada SD. Analysis of compositions of microbiomes with bias correction. Nat Commun. 2020;11.

35. Kriss M, Hazleton KZ, Nusbacher NM, Martin CG, Lozupone CA. Low diversity gut microbiota dysbiosis: drivers, functional implications and recovery. Curr Opin Microbiol. 2018;44:34–40.

36. Duvallet C, Gibbons SM, Gurry T, Irizarry RA, Alm EJ. Meta-analysis of gut microbiome studies identifies disease-specific and shared responses. Nat Commun. 2017;8.

37. Clemente C, Jose, Ursell K, Luke, Parfrey W, Laura, Knight R. The impact of the gut microbiota on human health: An integrative view. Cell. 2012;148:1258–70.

38. Arumugam M, Raes J, Pelletier E, Le Paslier D, Yamada T, Mende DR, et al. Enterotypes of the human gut microbiome. Nature. 2011;473:174–80.

39. Gibiino G, Lopetuso LR, Scaldaferri F, Rizzatti G, Binda C, Gasbarrini A. Exploring Bacteroidetes: Metabolic key points and immunological tricks of our gut commensals. Dig Liver Dis. 2018;50:635–9.

40. Spor A, Koren O, Ley R. Unravelling the effects of the environment and host genotype on the gut microbiome. Nat Rev Microbiol. 2011;9:279–90.

41. Feng J, Li B, Jiang X, Yang Y, Wells GF, Zhang T, et al. Antibiotic resistome in a large-scale healthy human gut microbiota deciphered by metagenomic and network analyses. Environ Microbiol. 2018;20:355–68.

42. Hu Y, Yang X, Qin J, Lu N, Cheng G, Wu N, et al. Metagenome-wide analysis of antibiotic resistance genes in a large cohort of human gut microbiota. Nat Commun. 2013;4.

43. Scallan E, Crim SM, Runkle A, Henao OL, Mahon BE, Hoekstra RM, et al. Bacterial enteric infections among older adults in the United States: Foodborne diseases active surveillance network, 1996-2012. Foodborne Pathog Dis. 2015;12:492–9.

44. Scallan E, Mahon BE, Hoekstra RM, Griffin PM. Estimates of illnesses, hospitalizations and deaths caused by major bacterial enteric pathogens in young children in the United States. Pediatr Infect Dis J. 2013;32:217–21.

45. Lupp C, Robertson ML, Wickham ME, Sekirov I, Champion OL, Gaynor EC, et al. Host-mediated inflammation disrupts the intestinal microbiota and promotes the overgrowth of Enterobacteriaceae. Cell Host Microbe. 2007;2:204.

46. Stecher B, Denzler R, Maier L, Bernet F, Sanders MJ, Pickard DJ, et al. Gut inflammation can boost horizontal gene transfer between pathogenic and commensal Enterobacteriaceae. Proc Natl Acad Sci. 2012;109:1269–74.

47. Skurnik D, Le Menac’H A, Zurakowski D, Mazel D, Courvalin P, Denamur E, et al. Integron-associated antibiotic resistance and phylogenetic grouping of *Escherichia coli* isolates from healthy subjects free of recent antibiotic exposure. Antimicrob Agents Chemother. 2005;49:3062–5.

48. Chow J, Tang H, Mazmanian SK. Pathobionts of the gastrointestinal microbiota and inflammatory disease. Curr Opin Immunol. 2011;23:473–80.

49. Wales A, Davies R. Co-selection of resistance to antibiotics, biocides and heavy metals, and its relevance to foodborne pathogens. Antibiotics. 2015;4:567–604.

50. Matchado MS, Lauber M, Reitmeier S, Kacprowski T, Baumbach J, Haller D, et al. Network analysis methods for studying microbial communities: A mini review. Comput Struct Biotechnol J. 2021;19:2687–98.

51. Shoemaker NB, Vlamakis H, Hayes K, Salyers AA. Evidence for extensive resistance gene transfer among Bacteroides spp. and among Bacteroides and other genera in the human colon. Appl Environ Microbiol. 2001;67:561–8.

52. Hedberg M, Nord CE, ESCMID Study Group on Antimicrobial Resistance in Anaerobic Bacteria. Antimicrobial susceptibility of Bacteroides fragilis group isolates in Europe. Clin Microbiol Infect. 2003;9:475–88.

53. Snydman DR, Jacobus NV, McDermott LA, Golan Y, Hecht DW, Goldstein EJ, et al. Lessons learned from the anaerobe survey: Historical perspective and review of the most recent data (2005-2007). Clin Infect Dis. 2010;50 Suppl 1:S26–33.

54. Teunis PFM, Evers EG, Hengeveld PD, Dierikx CM, Wielders CCCH, Van Duijkeren E. Time to acquire and lose carriership of ESBL/pAmpC producing *E. coli* in humans in the Netherlands. PLoS One. 2018;13:e0193834.

55. Doi Y, Adams-Haduch JM, Peleg AY, D’Agata EMC. The role of horizontal gene transfer in the dissemination of extended-spectrum beta-lactamase-producing *Escherichia coli* and *Klebsiella pneumoniae* isolates in an endemic setting. Diag Microbiol Infect Dis. 2012;74:34–8.

56. Whittle G, Shoemaker NB, Salyers AA. The role of *Bacteroides* conjugative transposons in the dissemination of antibiotic resistance genes. Cell Mol Life Sci. 2002;59:2044–54.

57. Shintani M, Sanchez ZK, Kimbara K. Genomics of microbial plasmids: Classification and identification based on replication and transfer systems and host taxonomy. 6; 2015.

58. Carr VR, Shkoporov A, Hill C, Mullany P, Moyes DL. Probing the mobilome: Discoveries in the dynamic microbiome. Trends Microbiol. 2021;29.

