## Supplemental tables and figures for "Recovery of the gut microbiome following enteric infection and persistence of antimicrobial resistance genes in specific microbial hosts"

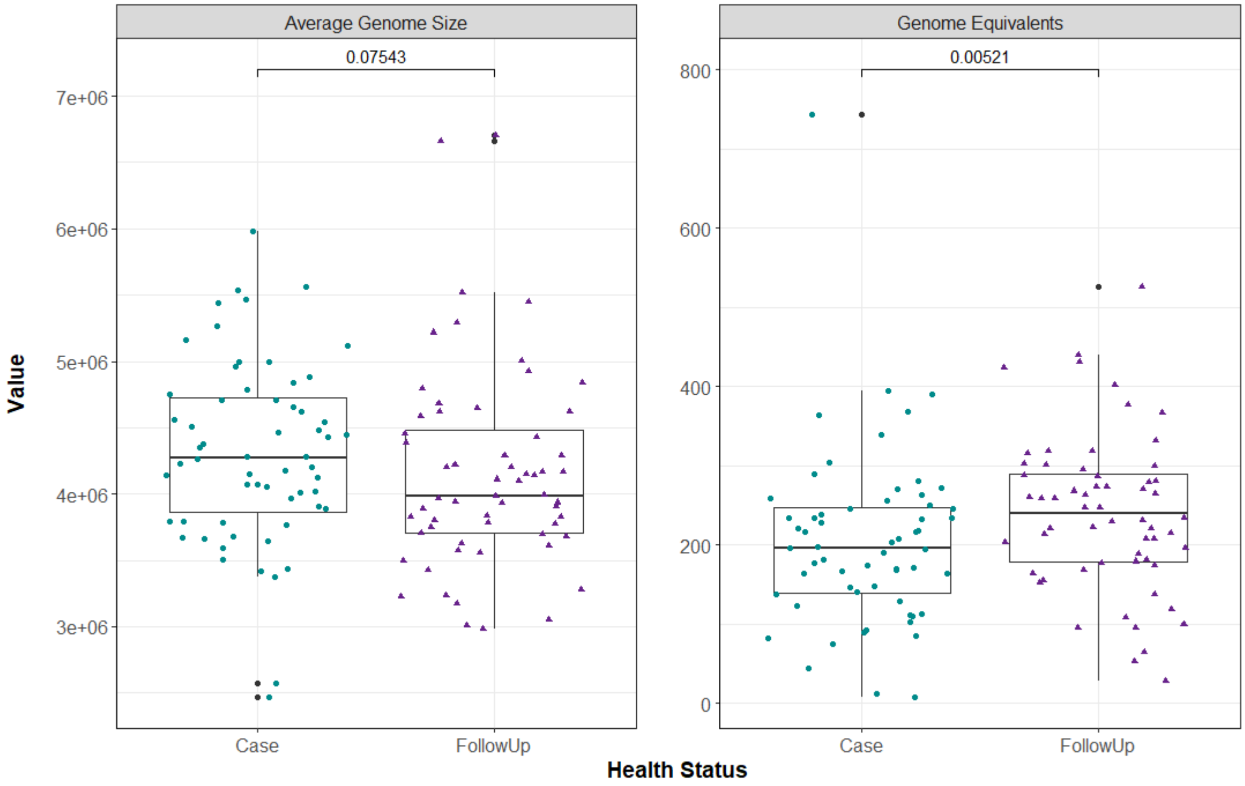


**Additional file 1. Average genome size (AGS) and estimated number of genome equivalents (GE) for the paired samples.** Each metric is displayed and stratified by health status. Case samples are represented by green circles, whereas the post-recovery samples (FollowUp) are shown as purple squares. Points are offset from the vertical to allow interpretation of all samples. The median of each measure is shown as a thick bar within the box and the first and third quartiles are represented by the bottom and top of the box, respectively. P-values were calculated using the Wilcoxon signed-rank test for paired samples and are shown above the comparison bar within each plot.

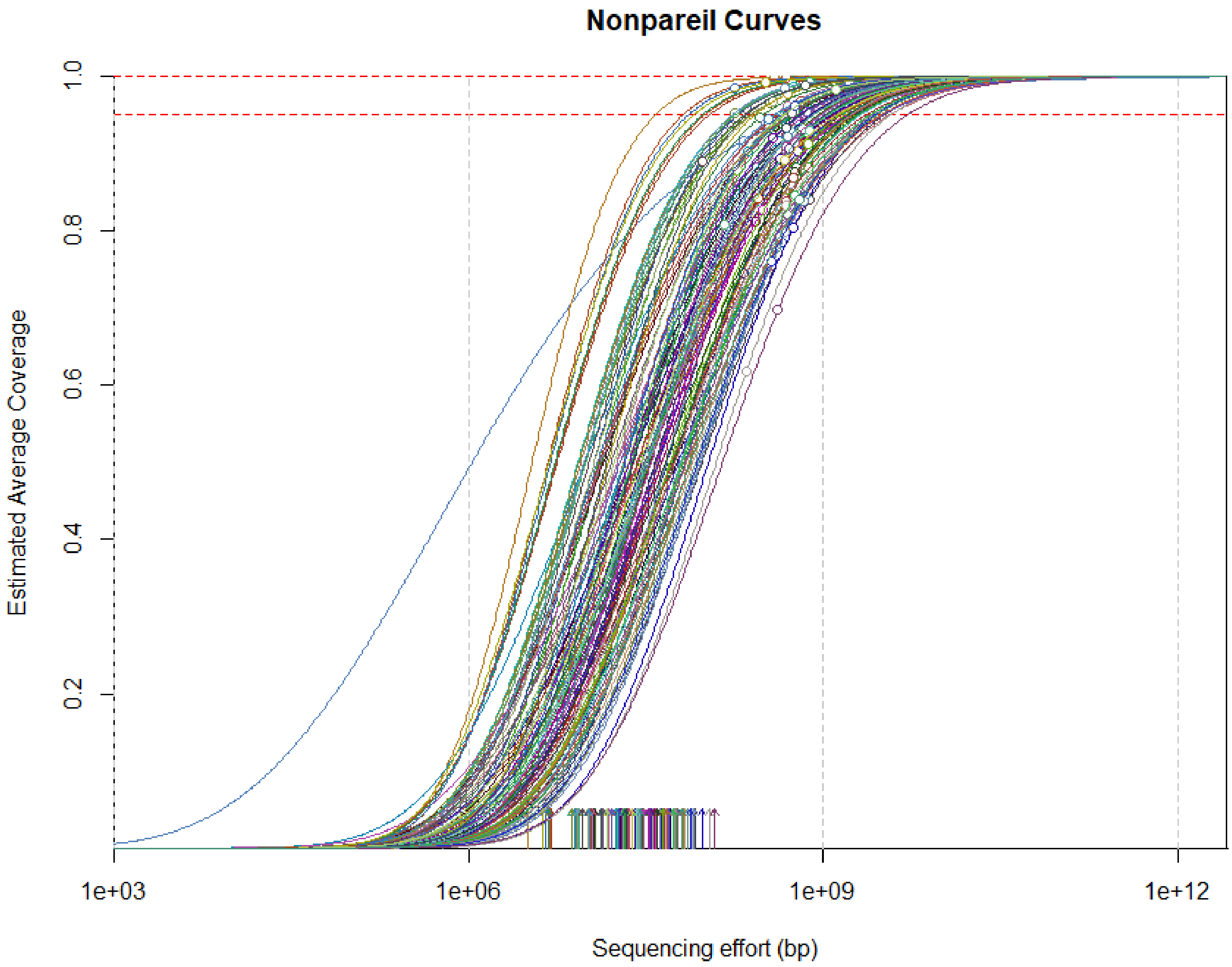


**Additional file 2. Metagenomic sequencing coverage of short paired-end reads was determined using Nonpareil.** The estimated sequencing coverage at increasing sequencing effort (s-curves) is shown as well as the actual coverage of each sample (open circles) for the case (n=60) and follow-up (n=60) samples. Each s-curve represents a single sample; arrows aligned on the x-axis represent the Nonpareil index of sequence diversity, a metric capturing the complexity of microbial communities in sequencing space. The box bordered by dotted red lines encapsulated coverage ranging from 95-100%. The overall mean coverage among cases and follow-ups was 86.3%, with a Nonpareil diversity score of 17.0. Additionally, the estimated sequencing effort among all cases and follow-ups was 4.77e+08 base pairs.

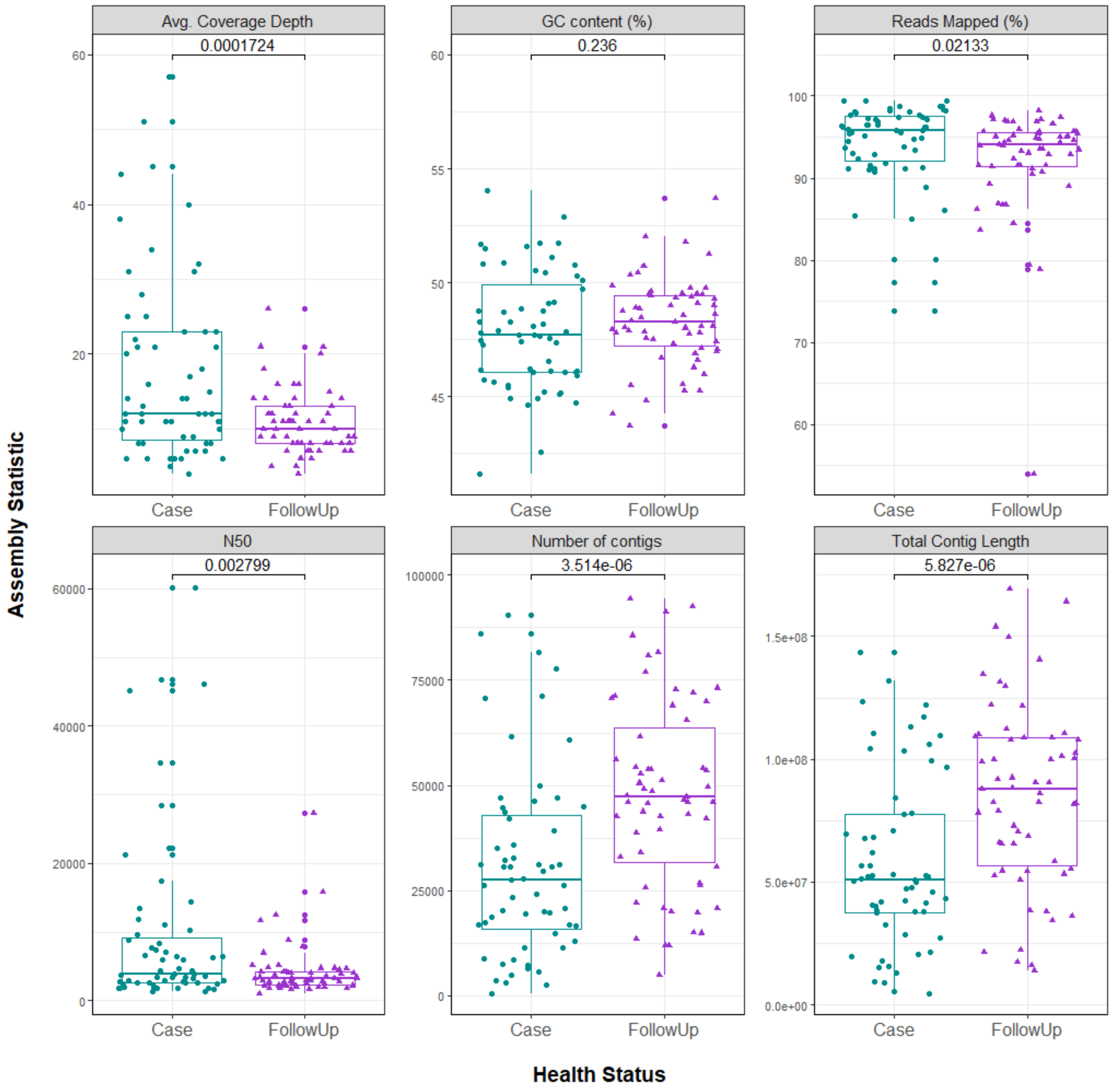


**Additional file 3. Assembly coverage statistics quantified by the Quality Assessment Tool for Genome Assemblies (QUAST).** Six different assembly statistics are shown stratified by health status, with samples represented by green circles (cases) or purple squares (follow-ups). Clockwise from the top-left, the six metrics include: average (Avg.) depth of coverage, percent of GC base pairs among samples, percentage of reads mapped to contigs, total contig length, number of contigs, and the N50 value. In each plot, points are offset from the vertical for clarity. The median is shown as a thick bar within the box (green for cases; purple for follow-ups) and the first and third quartiles are represented by the bottom and top of the box, respectively. P-values were calculated using the Wilcoxon signed-rank test for paired samples and are shown above the comparison bar within each plot.

**Additional file 4. Shapiro-Wilk Test results to determine whether the data for case and follow-up samples are normally distributed in the microbiome and resistome datasets.**

| Dataset | Alpha Diversity Metric | Test Statistic (W) | p-value |
|---|---|---|---|
| *Microbiome* | Richness | 0.936 | 2.35E-05 |
| | Shannon Diversity | 0.933 | 1.45E-05 |
| | Pielou's Evenness | 0.940 | 4.25E-05 |
| *Resistome* | Richness | 0.940 | 4.18E-05 |
| | Shannon Diversity | 0.905 | 3.63E-07 |
| | Pielou's Evenness | 0.885 | 3.68E-08 |

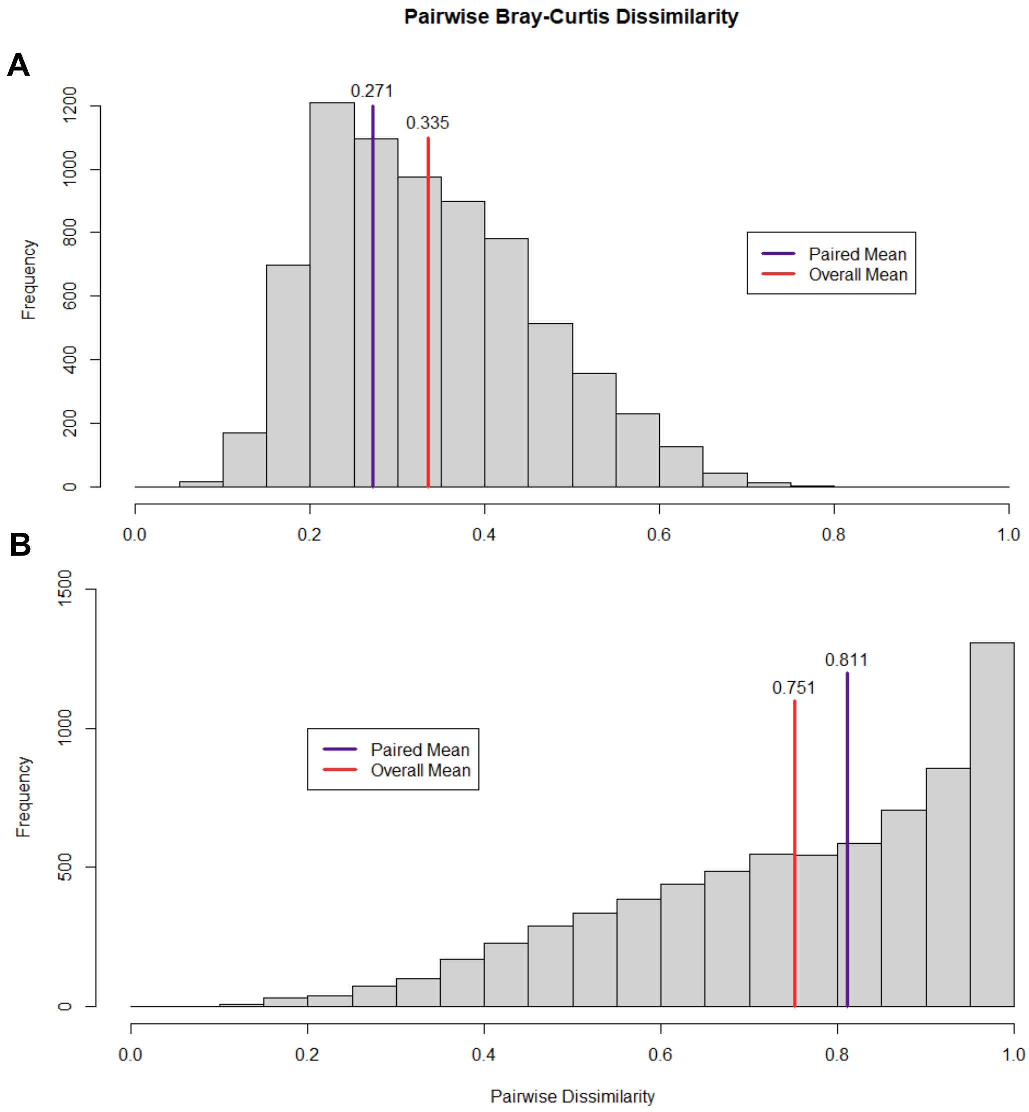


**Additional file 5. Pairwise comparison of Bray-Curtis dissimilarity among cases and follow-ups for the A) the microbiome (species-level) and B) resistome (gene-level) data**. Upon construction of the dissimilarity matrix, pairwise dissimilarity scores were output into two separate data frames: one contained all pairwise comparisons and one contained only pairwise comparisons among relevant paired samples. In each plot, the overall mean dissimilarity across all samples is displayed as a red line with the value of the mean oriented above the line (microbiome = 0.335; resistome = 0.751). The mean among paired samples only is shown as a purple line (microbiome = 0.271; resistome = 0.811). A Welch's t-test indicated that the dissimilarity among paired samples was significantly lower than the overall mean for microbiome composition (p = 2.58e-05; two-sided) and was significantly higher than the overall mean for the resistome (p = 0.013; two-sided).

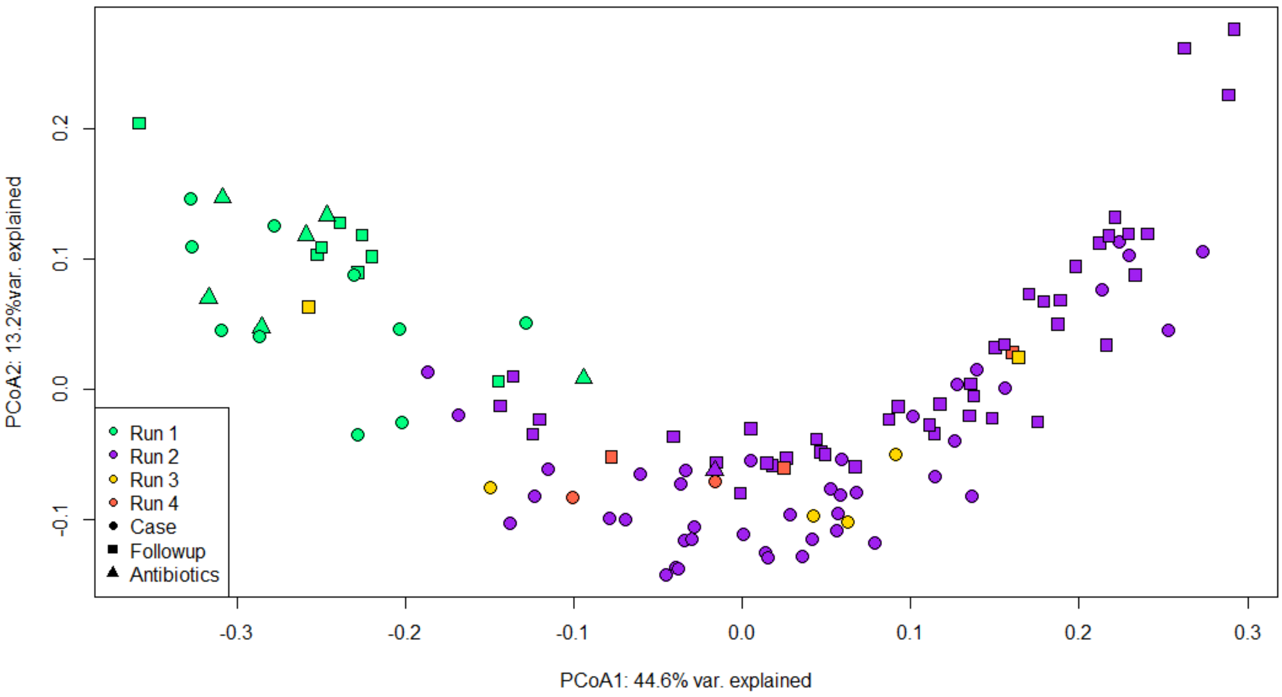


**Additional file 6. Exploring potential batch effects related to sequencing run using Principal Coordinate Analysis (PCoA) of cases and follow-ups.** The plot shows the distribution of cases (circles) and follow-ups (squares) based on Bray-Curtis dissimilarity calculated from species-level abundances of the gut microbiota. The first and second coordinate display the percentage of similarity explained. Points are colored by their corresponding sequencing run: Run 1 (green), Run 2 (purple), Run 3 (yellow), and Run 4 (orange). Samples sequenced in Run 1 cluster separately from those in Runs 2, 3, and 4 along the first axis. Patients that self-reported use of antibiotics two weeks prior to sample collection are indicated by triangles.

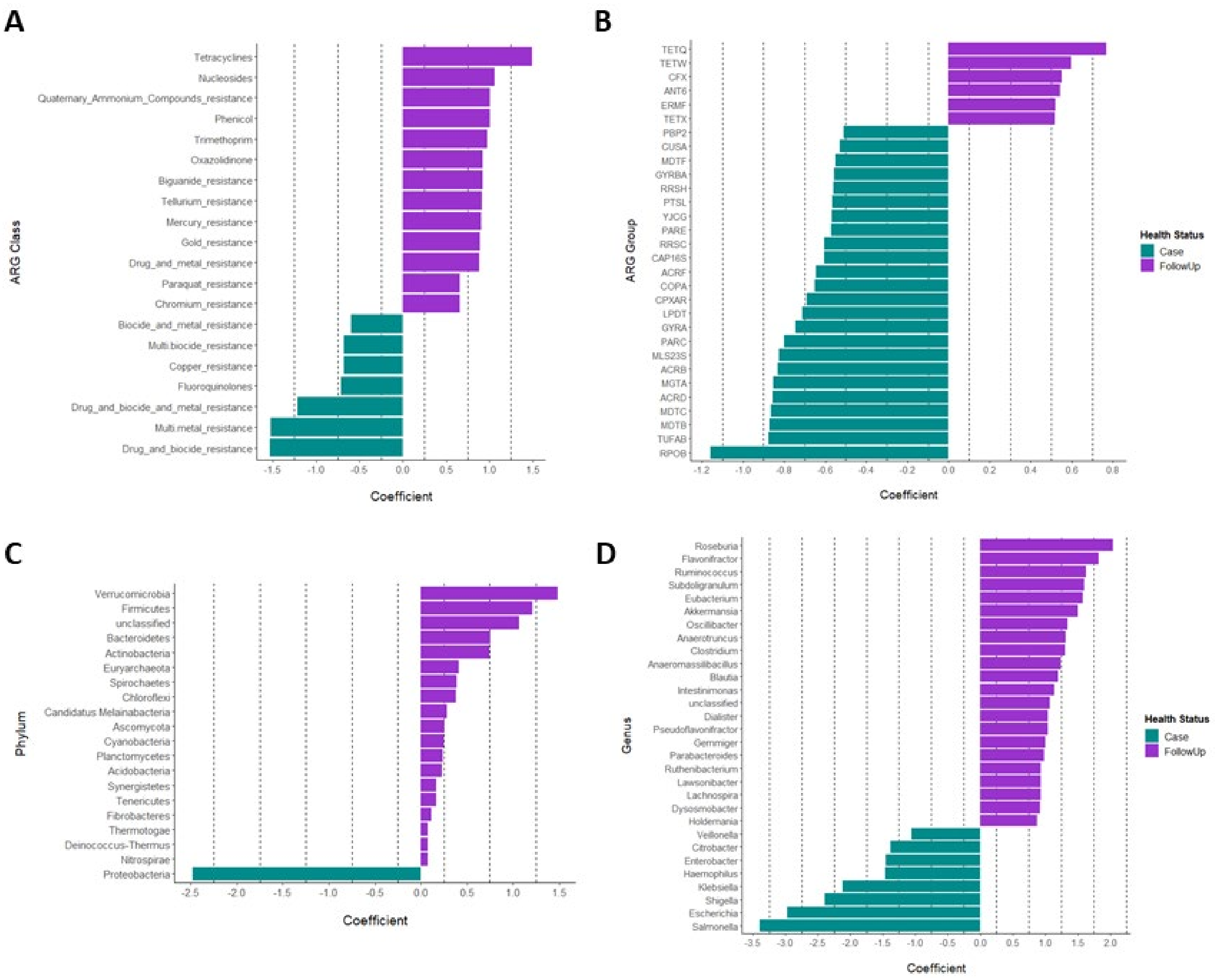


**Additional file 7. Differential abundance of taxonomic and resistance gene features for cases and follow-ups detected using ANCOM-BC.** Differentially abundant **A**) antimicrobial resistance gene (ARG) classes, **B**) ARG groups, **C**) taxonomic phyla, and **D**)

genera are shown. The x-axis displays the range of coefficients associated with each feature shown on the y-axis. In each sub-plot, purple bars designate resistome or microbiome features that were found to be more abundant in follow-ups, whereas teal bars show those features that were more abundant in cases.

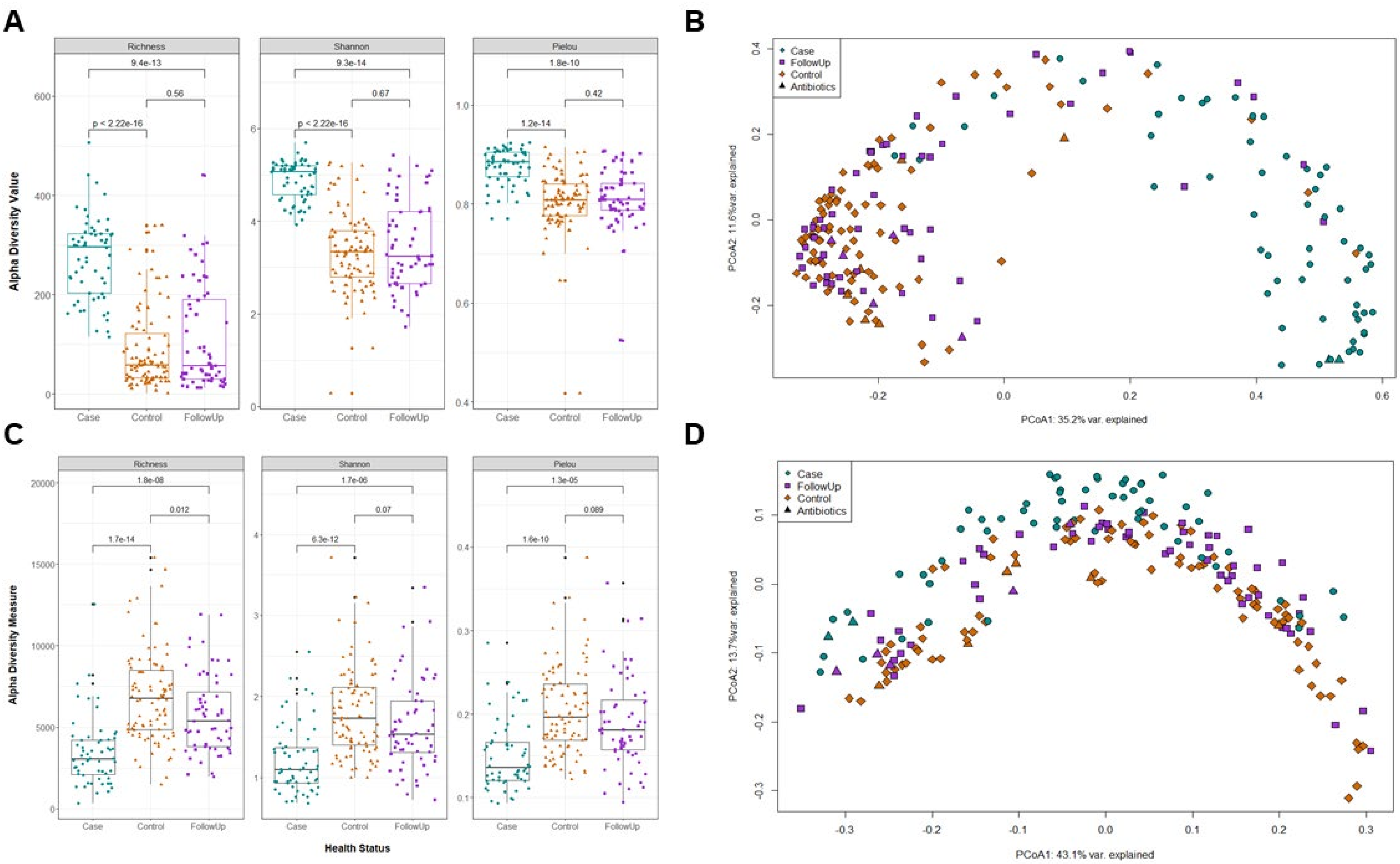

A
B
C
D
Richness
Shannon
Pielou
Alpha Diversity Value
Alpha Diversity Measure
Health Status
Case
Control
FollowUp
Antibiotics
9.4e-13
0.56
p < 2.22e-16
9.3e-14
0.67
1.8e-10
0.42
1.2e-14
1.8e-08
0.012
1.7e-14
1.7e-06
0.07
6.3e-12
1.3e-05
0.089
1.6e-10
PCoA1: 35.2% var. explained
PCoA2: 11.6%var. explained
PCoA1: 43.1% var. explained
PCoA2: 13.7%var. explained

**Additional file 8. Resistome and microbiome diversity among patients (Case), healthy household members (Control) and recovered cases (FollowUp).** Alpha diversity measures (Richness, Shannon's Diversity Index, and Pielou's Evenness Index) are shown for the **A)** resistome and **C)** microbiome stratified by health status. Samples indicated by green circles are the cases, whereas the purple squares represent the follow-ups, and the orange triangles are the household member controls. The median is indicated by the thick bar (green for cases; purple for follow-ups) and the first and third quartiles are shown at the bottom and top of the box, respectively. P-values were calculated using the Wilcoxon signed-rank test for paired samples and are indicated above the comparison bar within each plot. Beta-diversity is displayed via a Principal Coordinates Analysis (PCoA) plot of cases (green, circles), follow-ups (purple, squares), and controls (orange, diamonds) for the **B)** resistome and **D)** microbiome based on Bray-Curtis dissimilarity calculated from gene-level (resistome) or species-level (microbiome) abundances. The first and second coordinate are shown and include the corresponding percentage of similarity explained. Patients that self-reported use of antibiotics two weeks prior to sample collection are indicated by triangular data points.

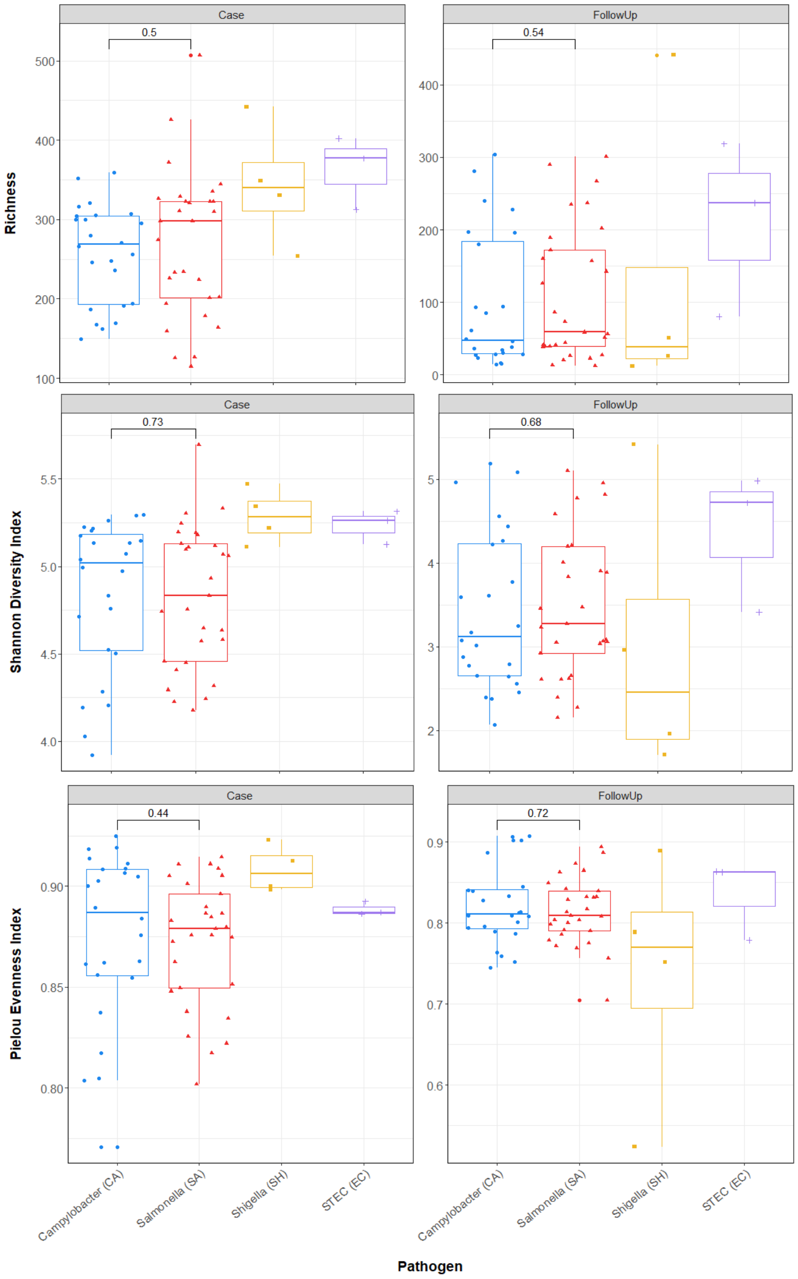
Case
FollowUp
0.5
0.54
Richness
0.73
0.68
Shannon Diversity Index
0.44
0.72
Pielou Evenness Index
Campylobacter (CA)
Salmonella (SA)
Shigella (SH)
STEC (EC)
Pathogen

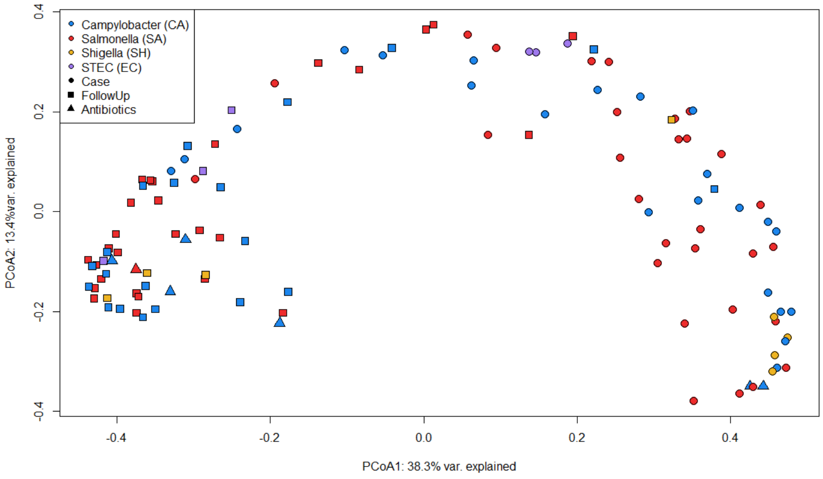
Campylobacter (CA)
Salmonella (SA)
Shigella (SH)
STEC (EC)
Case
FollowUp
Antibiotics
PCoA2: 13.4%var. explained
PCoA1: 38.3% var. explained

**Additional file 9. Alpha and beta diversity of the resistome do not differ across the four different enteric pathogens.** Box plots for the three resistome alpha diversity measures (Richness, Shannon's Diversity Index, and Pielou's Evenness Index) are shown with separate plots for the case (Case, left) and follow-up (FollowUp, right) samples. Each plot is stratified by the pathogen linked to the acute infection: *Campylobacter* (blue circles), *Salmonella* (red triangles), *Shigella* (yellow squares), and Shiga toxin-producing *Escherichia coli* (STEC; purple pluses). Points are offset from the vertical for clarity and the median is indicated by the thick bar. The first and third quartiles are represented by the bottom and top of the box, respectively. P-values comparing *Campylobacter* and *Salmonella* were calculated using the Wilcoxon rank-sum test and are shown above the comparison bar within each plot. Comparisons with S*higella* and STEC were not pursued because of low sample sizes. The bottom graph shows the Principal Coordinates Analysis (PCoA) plot based on Beta-diversity dissimilarity calculated from gene-level abundances for cases (circles) and follow-ups (squares). Patients reporting antibiotic use two weeks prior to sample collection are indicated by triangular data points. Colored points refer to the different pathogens: *Campylobacter* (blue), *Salmonella* (red), *Shigella* (yellow), STEC (purple). The first and second coordinate shows the corresponding percentage of similarity explained.

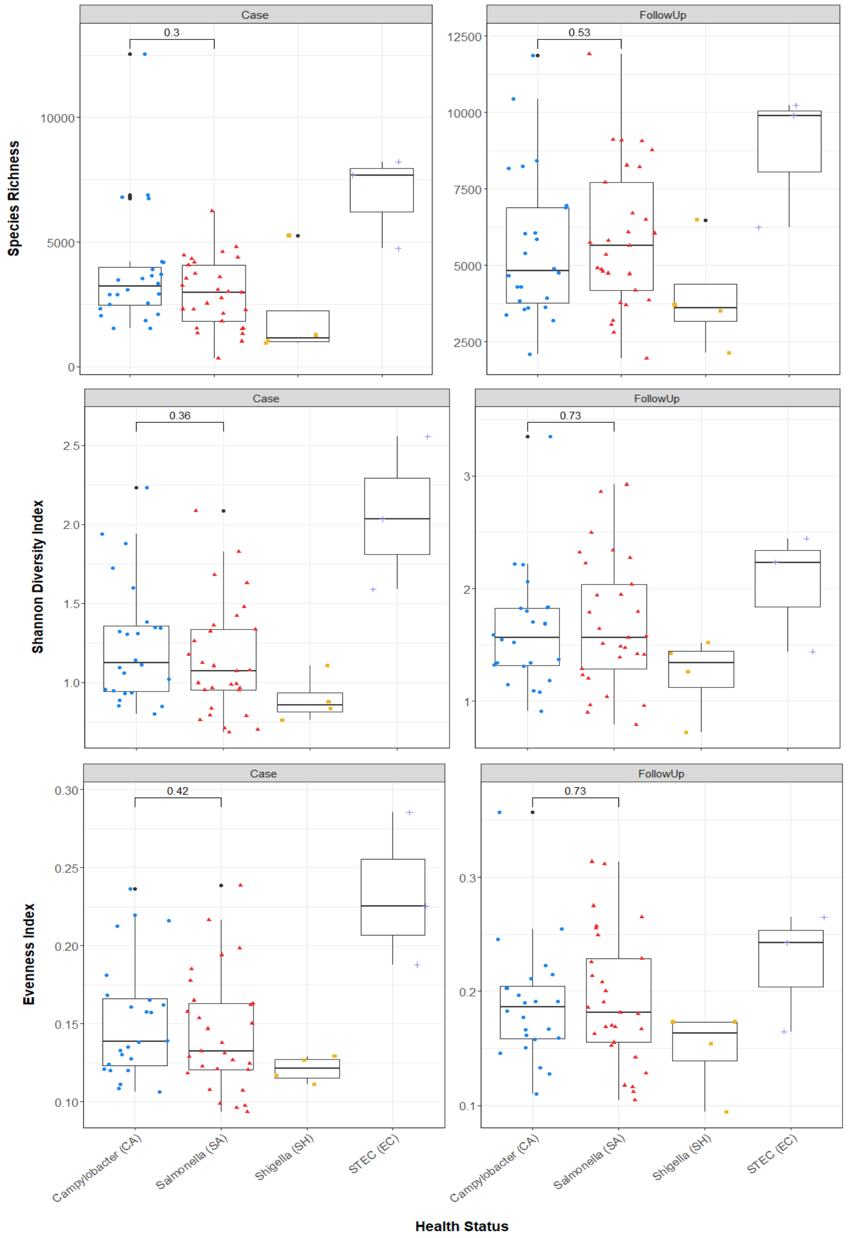

Case
FollowUp
Species Richness
Shannon Diversity Index
Evenness Index
Campylobacter (CA)
Salmonella (SA)
Shigella (SH)
STEC (EC)
Health Status


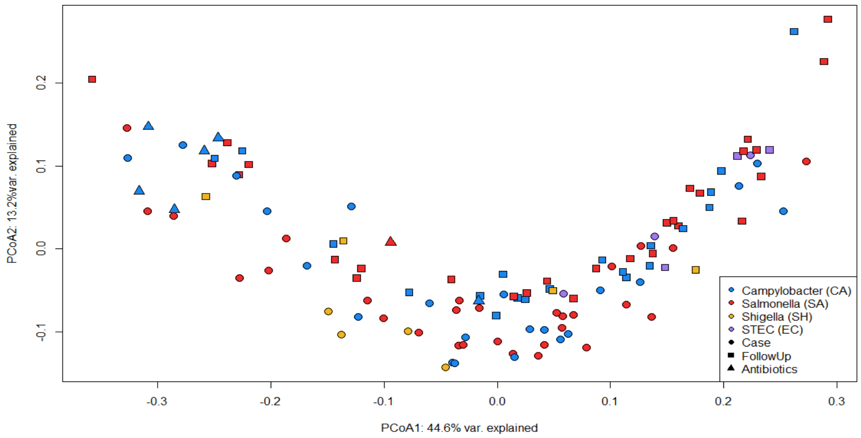

PCoA2: 13.2%var. explained
PCoA1: 44.6% var. explained
Campylobacter (CA)
Salmonella (SA)
Shigella (SH)
STEC (EC)
Case
FollowUp
Antibiotics

**Additional file 10. Microbiome diversity and composition do not differ across the four different enteric pathogens.** Box plots for the three microbiome alpha diversity measures (Richness, Shannon's Diversity Index, and Pielou's Evenness Index) are shown with separate plots for the case (Case, left) and follow-up (FollowUp, right) samples. Each plot is stratified by the pathogen linked to the acute infection: *Campylobacter* (blue circles), *Salmonella* (red triangles), *Shigella* (yellow squares), and Shiga toxin-producing *Escherichia coli* (STEC; purple pluses). Points are offset from the vertical for clarity and the median is indicated by the thick bar. The first and third quartiles are represented by the bottom and top of the box, respectively. P-values comparing *Campylobacter* and *Salmonella* were calculated using the Wilcoxon rank-sum test and are shown above the comparison bar within each plot. The bottom panel shows the Principal Coordinates Analysis plot based on Beta-diversity dissimilarity calculated from gene-level abundances for cases (circles) and follow-ups (squares). Points are coded for antibiotic use (triangles) and the four different pathogens: *Campylobacter* (blue), *Salmonella* (red), *Shigella* (yellow), STEC (purple). The first and second coordinate shows the corresponding percentage of similarity explained.

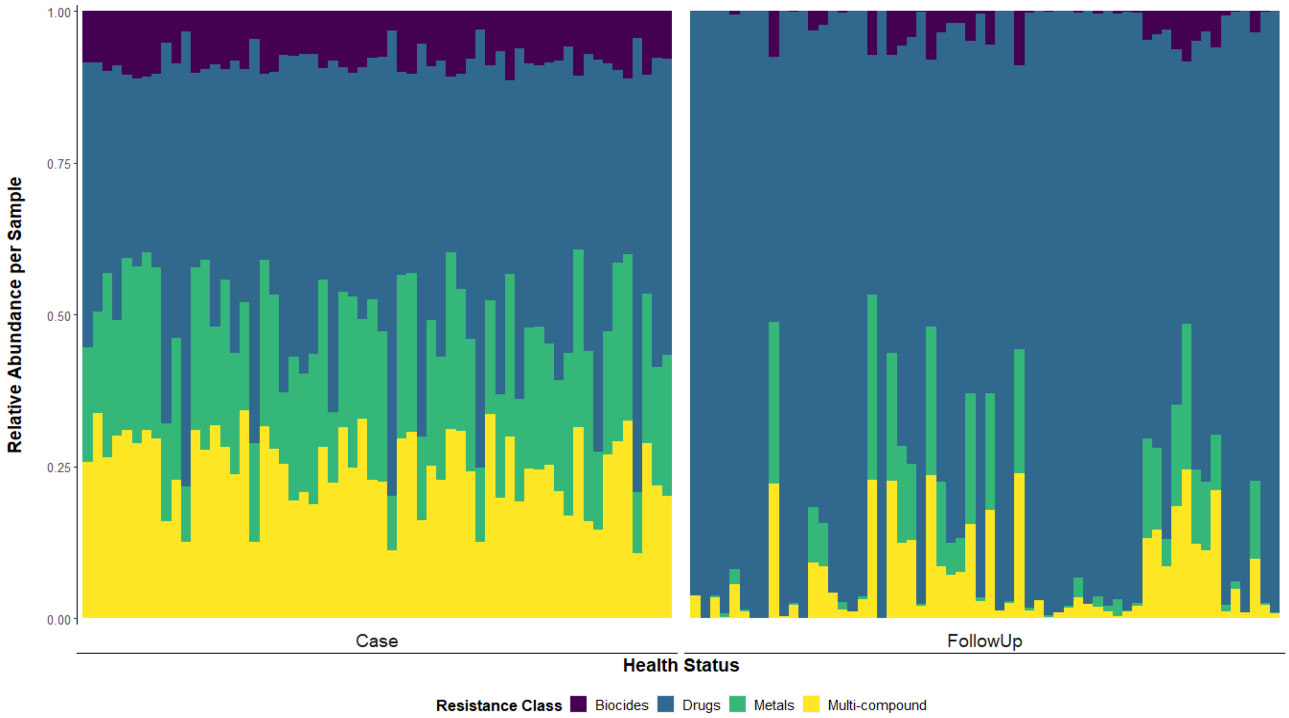


**Additional file 11. Relative abundance of resistance gene types in case and follow-up samples.** The relative abundance of resistance genes assigned to biocides, drugs, metals, and multi-compounds is shown for cases (Case, left panel) and follow-ups (FollowUp, right panel), with each column representing the resistome from one individual. Columns are ordered by the paired sample, meaning that the column position in each side of the plot refers to the same individual either during or after enteric infection. Relative abundances were determined using raw gene abundances that had been normalized by the estimated number of genome equivalents in a given sample.

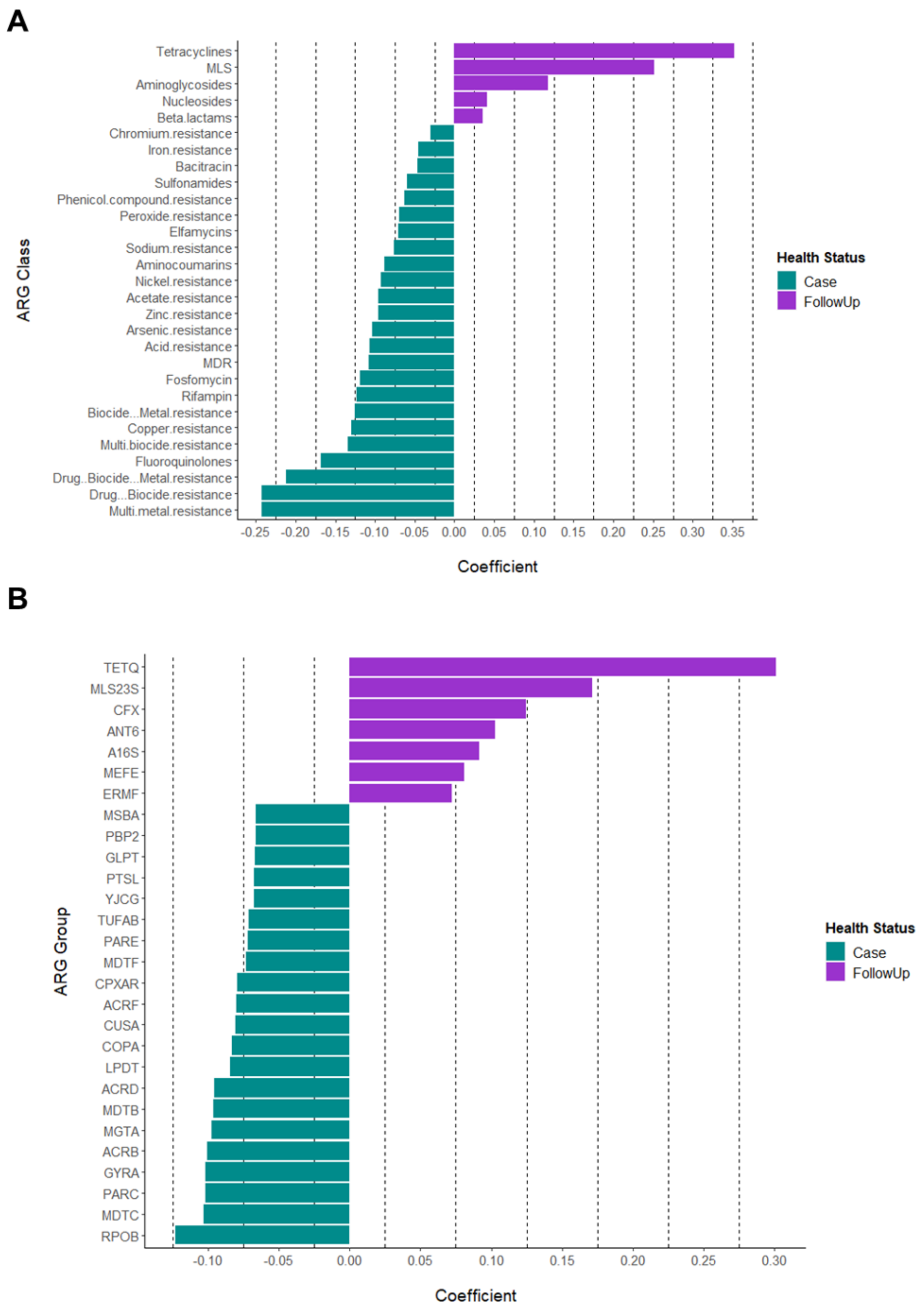


**Additional file 12. Differentially abundant ARG classes and groups among cases and follow-ups.** MMUPHin was used to identify ARG features of differential abundance. Coefficients for each ARG class (A) or ARG group (B) are shown on the x-axis with a cutoff of absolute value = 0.05; positive coefficients indicate ARG features which were more abundant among follow-ups, while those with negative coefficients were more prominent in cases. The specific ARG features are displayed on the y-axis. The bars in the plot are colored by health status with which that particular feature is associated (cases=green, follow-ups=purple).

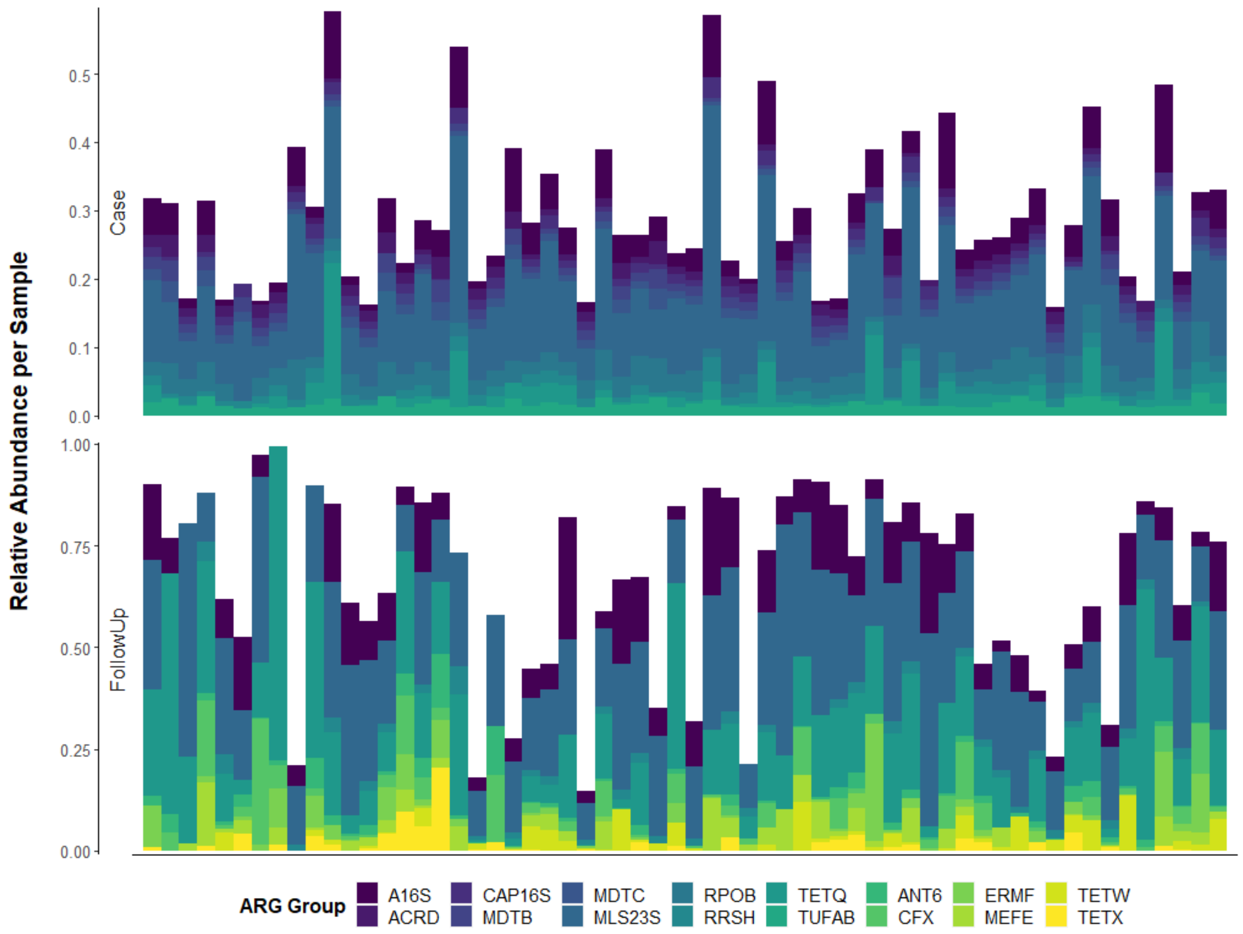


**Additional file 13. Relative abundance of the top-10 resistance gene groups among cases and follow-ups.** The relative abundance of resistance genes assigned to the top-10 most abundant groups is shown for cases (Case, top panel) and follow-ups (FollowUp, bottom panel), with each column representing the resistome from one individual. Columns are ordered by individual, meaning that the column position in each side of the plot refers to the samples from the same individual during (top) or after (bottom) enteric infection. Relative abundances were determined using raw gene abundances that had been normalized by the estimated number of genome equivalents in a sample.

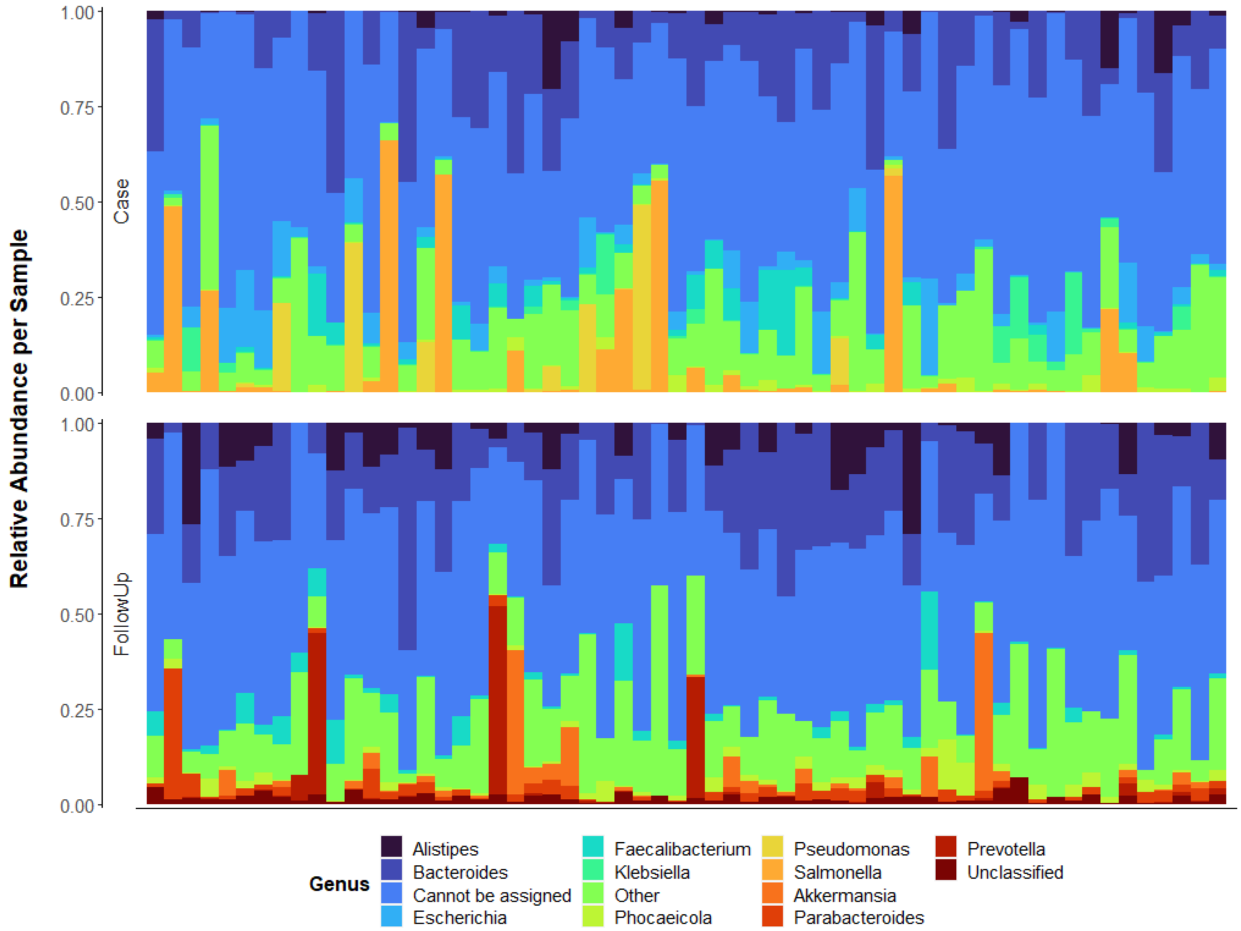


**Additional file 14. Relative abundance of microbial genera differ between cases and follow-ups.** The top-10 microbial genus with the greatest average relative abundance among cases or follow-ups is shown with each column representing the microbiome from one individual. Columns are ordered by their sample pairing, meaning that the column position for each facet of the plot refers to the same individual either during (Case; Top) or after (FollowUp; Bottom) enteric infection. Relative abundances were determined using raw gene abundances that had been normalized by the approximate number of genome equivalents in the sample as determined using MicrobeCensus

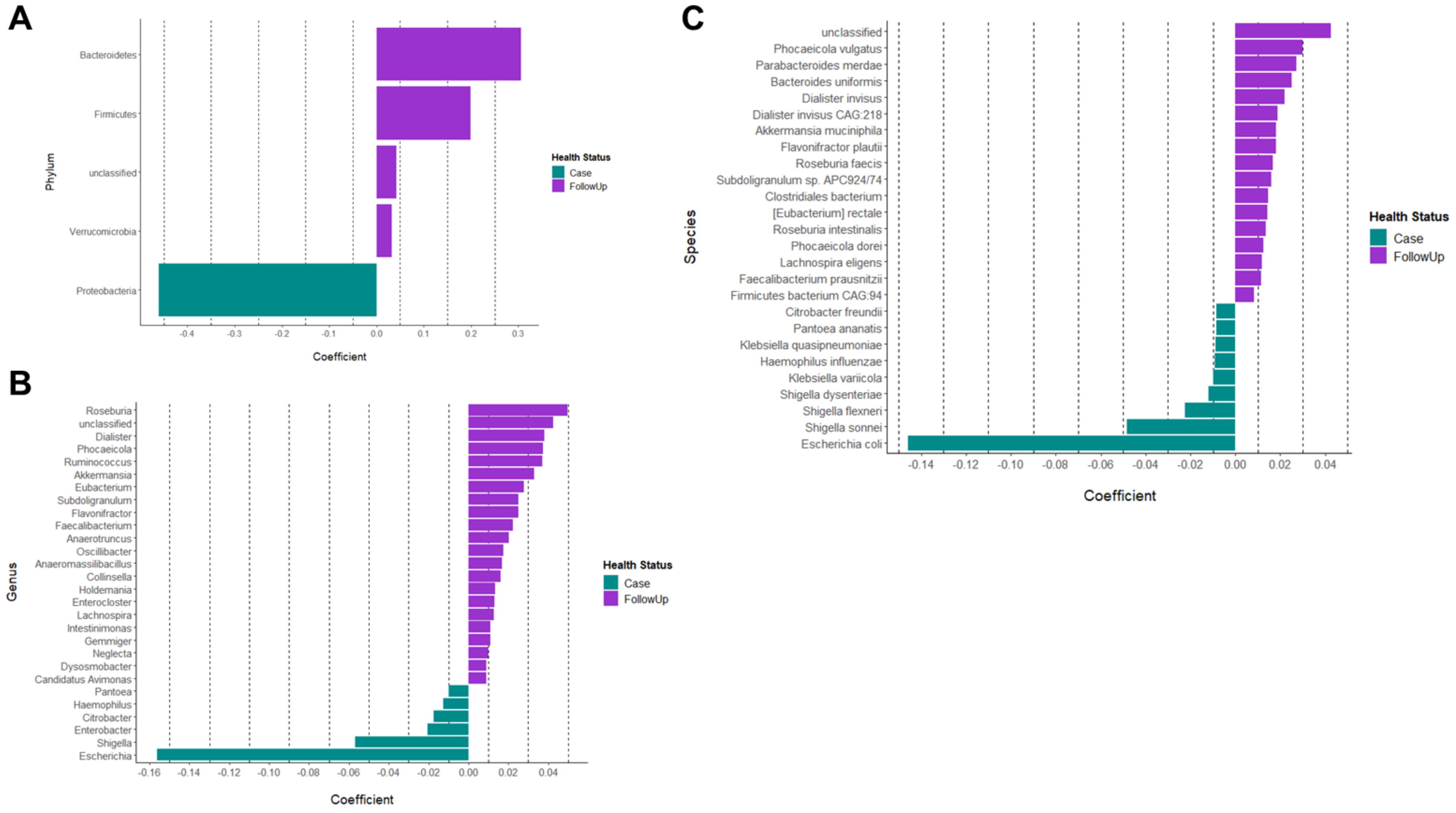


**Additional file 15. Differential abundance of phyla, genera, and species among cases and follow-ups.** MMUPHin was used to identify microbial features of differential abundance. Coefficients for each phylum (A) genus (B) or species (C) are shown on the x-axis with a cutoff of absolute value=0.05, 0.008, and 0.008, respectively; positive coefficients indicate features that were more abundant among follow-ups, while negative coefficients indicate those that were more abundant cases. The specific taxonomic features are displayed on the y-axis. Bars in the plot are colored by sample (cases=green, follow-ups=purple).

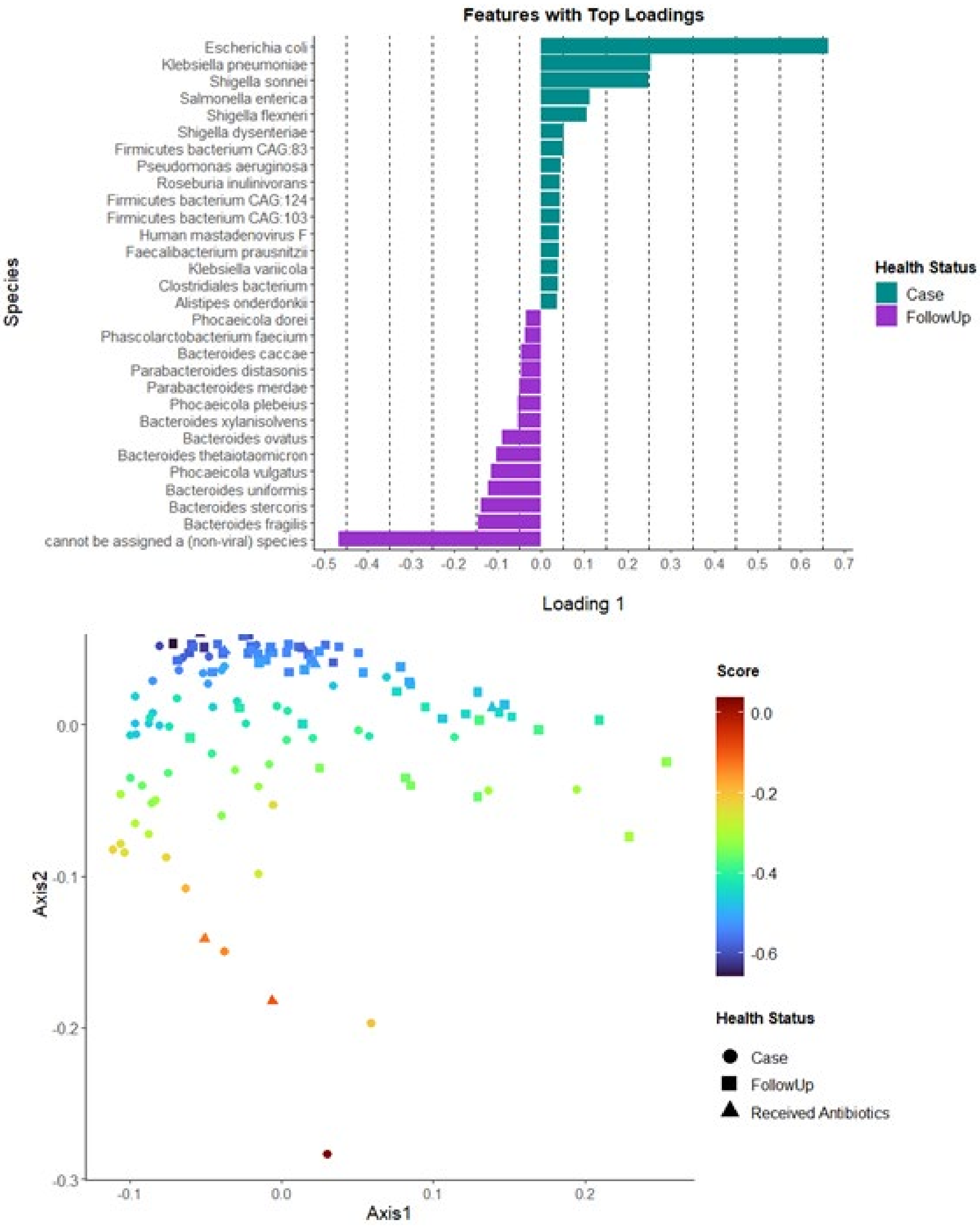


**Additional file 16. Continuous structure analysis reveals species gradients among cases and follow-ups.** MMUPHin was used to investigate potential continuous structure among the microbiome composition of cases and follow-ups at the species level. **A)** Species determined to comprise the top consensus loadings are shown; colors have been assigned to the loadings based on the sample affiliated with each loading (drawn from differential abundance analyses). Cases

(Case) are indicated in green and follow-ups (FollowUp) in purple. **B)** The species composition gradient is shown overlaid onto an ordination plot based on Bray-Curtis dissimilarity of case and follow-up microbiomes at the species level. Cases (circles), follow-ups (squares), and individuals who received antibiotics (triangles) are shown. The color gradient ("Score") refers to the continuous structure score affiliated with Loading 1. Juxtaposition of (A) and (B) allow interpretation of species tradeoffs that occur within the sample set (e.g., many cases contain higher levels of *Escherichia coli* at the expense of more beneficial bacteria such as Bacteroides species associated with the opposite direction.

**Additional file 17. Summary of beta-lactamase genes and their corresponding microbial hosts in cases and follow-ups.** The number of resistance genes in cases vs. follow-ups is shown and genes encoding extended-spectrum beta-lactamase (ESBL) production are indicted with an asterisk (*).

| Microbial Host | Beta-lactamase | Cases | Follow-ups |
|---|---|---|---|
| ***Enterobacter*** | ACT family cephalosporin-hydrolyzing class C beta-lactamase | 5 | 0 |
| ***Acinetobacter*** | ADC family extended-spectrum class C beta-lactamase* | 2 | 0 |
| ***Klebsiella*** | Beta-lactamase | 0 | 1 |
| **Uncultured** | beta-lactamase | 0 | 1 |
| ***Klebsiella*** | beta-lactamase | 4 | 0 |
| ***Escherichia*** | beta-lactamase | 3 | 6 |
| ***Escherichia*** | beta-lactamase TEM-1 variant* | 1 | 0 |
| **Synthetic** | beta-lactamase TEM-1 variant* | 19 | 4 |
| ***Klebsiella*** | beta-lactamase, partial | 3 | 0 |
| ***Shigella*** | BlaEC family class C beta-lactamase | 1 | 0 |
| ***Escherichia*** | BlaEC family class C beta-lactamase | 49 | 19 |
| ***Bacteroides*** | CepA family class A extended-spectrum beta-lactamase* | 19 | 13 |
| ***Prevotella*** | CfxA family class A broad-spectrum beta-lactamase | 5 | 9 |
| ***Bacteroides*** | CfxA family class A broad-spectrum beta-lactamase | 46 | 48 |
| ***Atlantibacter*** | class A beta-lactamase | 1 | 0 |
| ***Burkholderia*** | class A beta-lactamase | 0 | 1 |
| ***Proteus*** | class A beta-lactamase | 1 | 0 |
| ***Salmonella*** | class A beta-lactamase | 1 | 0 |
| ***Yersinia*** | class A beta-lactamase | 0 | 1 |
| ***Bacillus*** | class A beta-lactamase | 1 | 1 |
| ***Clostridium*** | class A beta-lactamase | 1 | 3 |
| ***Klebsiella*** | class A beta-lactamase, partial | 1 | 0 |
| ***Escherichia*** | class A beta-lactamase, partial | 0 | 2 |
| ***Bacteroides*** | class A beta-lactamase, subclass A2 | 45 | 47 |
| ***Providencia*** | class C beta-lactamase | 1 | 0 |
| ***Hafnia*** | class C beta-lactamase | 3 | 0 |
| ***Pseudomonas*** | class C beta-lactamase | 7 | 0 |
| ***Flavobacterium*** | class D beta-lactamase | 4 | 6 |

| Microbial Host | Beta-lactamase | Cases | Follow-ups |
|---|---|---|---|
| ***Klebsiella*** | class D beta-lactamase, partial | 1 | 0 |
| ***Salmonella*** | CMY family class C beta-lactamase | 3 | 2 |
| ***Salmonella*** | CMY-2 family class C beta-lactamase | 2 | 1 |
| ***Citrobacter*** | CMY-2 family class C beta-lactamase | 8 | 3 |
| ***Morganella*** | DHA family class C beta-lactamase | 1 | 0 |
| ***Klebsiella*** | LEN family class A beta-lactamase | 1 | 2 |
| ***Enterobacter*** | MIR family cephalosporin-hydrolyzing class C beta-lactamase | 1 | 0 |
| ***Klebsiella*** | MULTISPECIES: OXY family class A extended-spectrum beta-lactamase* | 2 | 0 |
| ***Klebsiella*** | OXA-1 family class D beta-lactamase* | 2 | 0 |
| ***Pseudomonas*** | OXA-50 family oxacillin-hydrolyzing class D beta-lactamase* | 2 | 0 |
| ***Acinetobacter*** | OXA-51 family carbapenem-hydrolyzing class D beta-lactamase* | 2 | 0 |
| ***Campylobacter*** | OXA-61 family class D beta-lactamase* | 2 | 0 |
| ***Klebsiella*** | OXY family class A extended-spectrum beta-lactamase* | 2 | 0 |
| ***Pseudomonas*** | PDC family class C beta-lactamase | 2 | 0 |
| ***Burkholderia*** | PenA family class A beta-lactamase | 1 | 0 |
| ***Raoultella*** | PLA/ORN/TER family class A beta-lactamase | 2 | 0 |
| ***Citrobacter*** | SED family class A beta-lactamase | 4 | 3 |
| ***Klebsiella*** | SHV family class A beta-lactamase* | 8 | 2 |
| ***Desulfovibrio*** | subclass B1 metallo-beta-lactamase | 0 | 1 |
| ***Myxococcus*** | subclass B1 metallo-beta-lactamase | 1 | 0 |
| ***Bacteroides*** | subclass B1 metallo-beta-lactamase | 8 | 14 |

**Additional file 18. Beta-lactamase genes and their corresponding microbial hosts identified in paired case and follow-up samples.** The number of resistance genes "lost" (i.e., present in the case sample but absent in the follow-up sample), "maintained" (i.e., present in both samples), or "acquired" (absent in case sample, present in follow-up sample) is shown. Genes encoding extended-spectrum beta-lactamase (ESBL) production are indicted with an asterisk (*).

| Microbial Host | Beta-lactamase | Lost | Maintained | Acquired |
|---|---|---|---|---|
| ***Enterobacter*** | ACT family cephalosporin-hydrolyzing class C beta-lactamase | 5 | 0 | 0 |
| ***Acinetobacter*** | ADC family extended-spectrum class C beta-lactamase* | 2 | 0 | 0 |
| ***Klebsiella*** | Beta-lactamase | 0 | 0 | 1 |
| **Uncultured** | beta-lactamase | 0 | 0 | 1 |
| ***Klebsiella*** | beta-lactamase | 4 | 0 | 0 |
| ***Escherichia*** | beta-lactamase | 3 | 0 | 6 |
| ***Escherichia*** | beta-lactamase TEM-1 variant* | 1 | 0 | 0 |
| **Synthetic** | beta-lactamase TEM-1 variant* | 18 | 1 | 3 |
| ***Klebsiella*** | beta-lactamase, partial | 3 | 0 | 0 |
| ***Shigella*** | BlaEC family class C beta-lactamase | 1 | 0 | 0 |
| ***Escherichia*** | BlaEC family class C beta-lactamase | 35 | 14 | 5 |
| ***Bacteroides*** | CepA family class A extended-spectrum beta-lactamase* | 9 | 10 | 3 |
| ***Prevotella*** | CfxA family class A broad-spectrum beta-lactamase | 3 | 2 | 7 |
| ***Bacteroides*** | CfxA family class A broad-spectrum beta-lactamase | 7 | 39 | 9 |
| ***Atlantibacter*** | class A beta-lactamase | 1 | 0 | 0 |
| ***Burkholderia*** | class A beta-lactamase | 0 | 0 | 1 |
| ***Proteus*** | class A beta-lactamase | 1 | 0 | 0 |
| ***Salmonella*** | class A beta-lactamase | 1 | 0 | 0 |
| ***Yersinia*** | class A beta-lactamase | 0 | 0 | 1 |
| ***Bacillus*** | class A beta-lactamase | 1 | 0 | 1 |
| ***Clostridium*** | class A beta-lactamase | 1 | 0 | 3 |
| ***Klebsiella*** | class A beta-lactamase, partial | 1 | 0 | 0 |
| ***Escherichia*** | class A beta-lactamase, partial | 0 | 0 | 2 |
| ***Bacteroides*** | class A beta-lactamase, subclass A2 | 7 | 38 | 9 |
| ***Providencia*** | class C beta-lactamase | 1 | 0 | 0 |
| ***Hafnia*** | class C beta-lactamase | 3 | 0 | 0 |

| Microbial Host | Beta-lactamase | Lost | Maintained | Acquired |
|---|---|---|---|---|
| ***Pseudomonas*** | class C beta-lactamase | 7 | 0 | 0 |
| ***Flavobacterium*** | class D beta-lactamase | 0 | 4 | 2 |
| ***Klebsiella*** | class D beta-lactamase, partial | 1 | 0 | 0 |
| ***Salmonella*** | CMY family class C beta-lactamase | 3 | 0 | 2 |
| ***Salmonella*** | CMY-2 family class C beta-lactamase | 2 | 0 | 1 |
| ***Citrobacter*** | CMY-2 family class C beta-lactamase | 6 | 2 | 1 |
| ***Morganella*** | DHA family class C beta-lactamase | 1 | 0 | 0 |
| ***Klebsiella*** | LEN family class A beta-lactamase | 1 | 0 | 2 |
| ***Enterobacter*** | MIR family cephalosporin-hydrolyzing class C beta-lactamase | 1 | 0 | 0 |
| ***Klebsiella*** | MULTISPECIES: OXY family class A extended-spectrum beta-lactamase* | 2 | 0 | 0 |
| ***Klebsiella*** | OXA-1 family class D beta-lactamase* | 2 | 0 | 0 |
| ***Pseudomonas*** | OXA-50 family oxacillin-hydrolyzing class D beta-lactamase* | 2 | 0 | 0 |
| ***Acinetobacter*** | OXA-51 family carbapenem-hydrolyzing class D beta-lactamase* | 2 | 0 | 0 |
| ***Campylobacter*** | OXA-61 family class D beta-lactamase* | 2 | 0 | 0 |
| ***Klebsiella*** | OXY family class A extended-spectrum beta-lactamase* | 2 | 0 | 0 |
| ***Pseudomonas*** | PDC family class C beta-lactamase | 2 | 0 | 0 |
| ***Burkholderia*** | PenA family class A beta-lactamase | 1 | 0 | 0 |
| ***Raoultella*** | PLA/ORN/TER family class A beta-lactamase | 2 | 0 | 0 |
| ***Citrobacter*** | SED family class A beta-lactamase | 4 | 0 | 3 |
| ***Klebsiella*** | SHV family class A beta-lactamase* | 8 | 0 | 2 |
| ***Desulfovibrio*** | subclass B1 metallo-beta-lactamase | 0 | 0 | 1 |
| ***Myxococcus*** | subclass B1 metallo-beta-lactamase | 1 | 0 | 0 |
| ***Bacteroides*** | subclass B1 metallo-beta-lactamase | 3 | 5 | 9 |

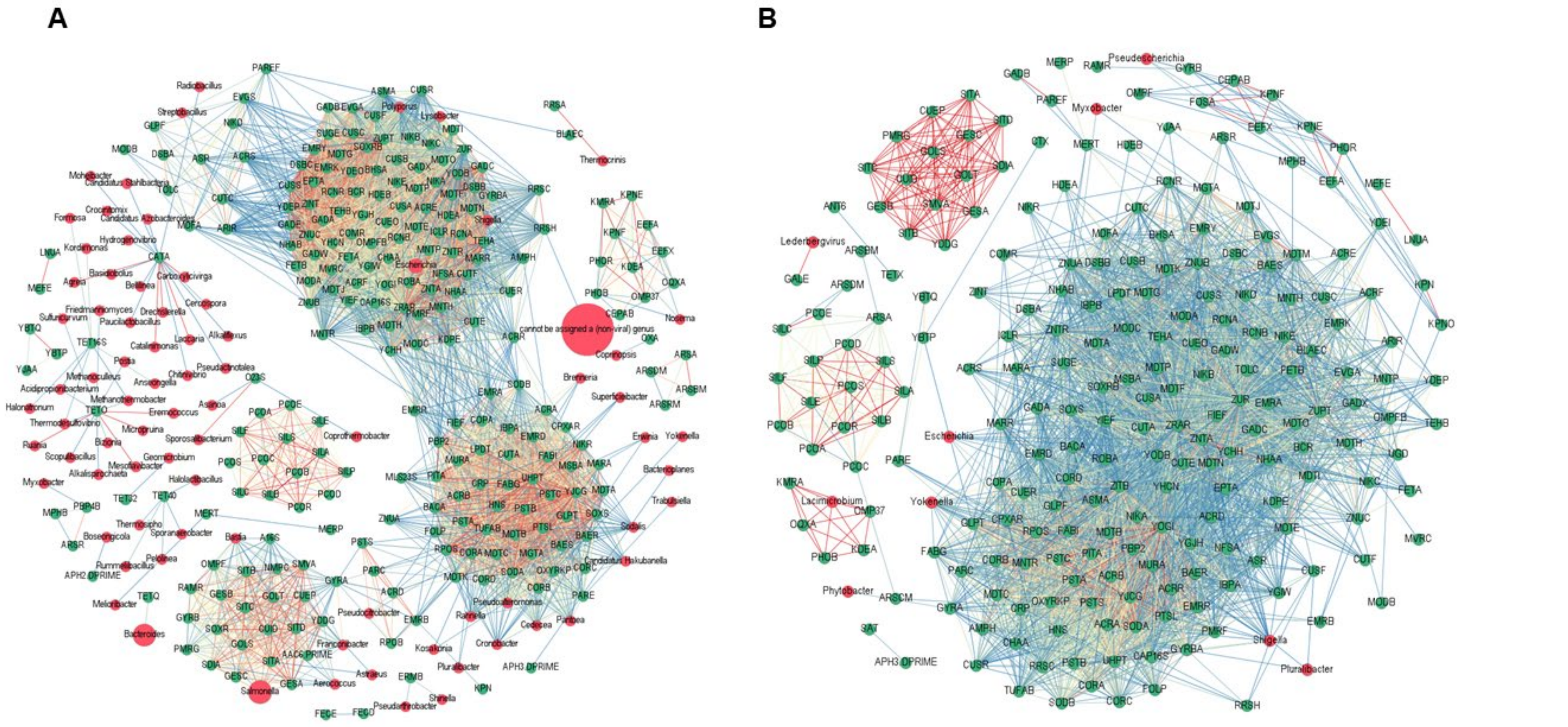


**Additional file 19**. **ARG-ARG and ARG-taxa connections are different for individuals infected with and recovering from *Salmonella*.** Correlation co-occurrence networks were constructed in Gephi 0.9.2 using Spearman's Rank correlation coefficients generated with the R-package 'Hmisc' (v4.5-0) for cases infected with *Salmonella* **(A)** or recovering (follow-ups) **(B)**. These networks display all ARG-ARG and ARG-taxa connections; taxa-taxa connections were excluded for clarity. Correlations included in the network all passed a cutoff of ρ>0.80 and q-value < 0.05. Nodes are colored by their identity as a taxonomic genus (red) or ARG group (green). Nodes are sized based on their overall abundance among samples; the larger the node, the more abundant. Nodes with ≥ 1 connection were included (i.e. degree cutoff=1). The edge color displays the strength of correlation, with blue demonstrating relatively weaker correlations (yet still >0.80), yellow showing medium correlation, and red showing strong correlation. Nodes are labeled with their corresponding genus or ARG group.

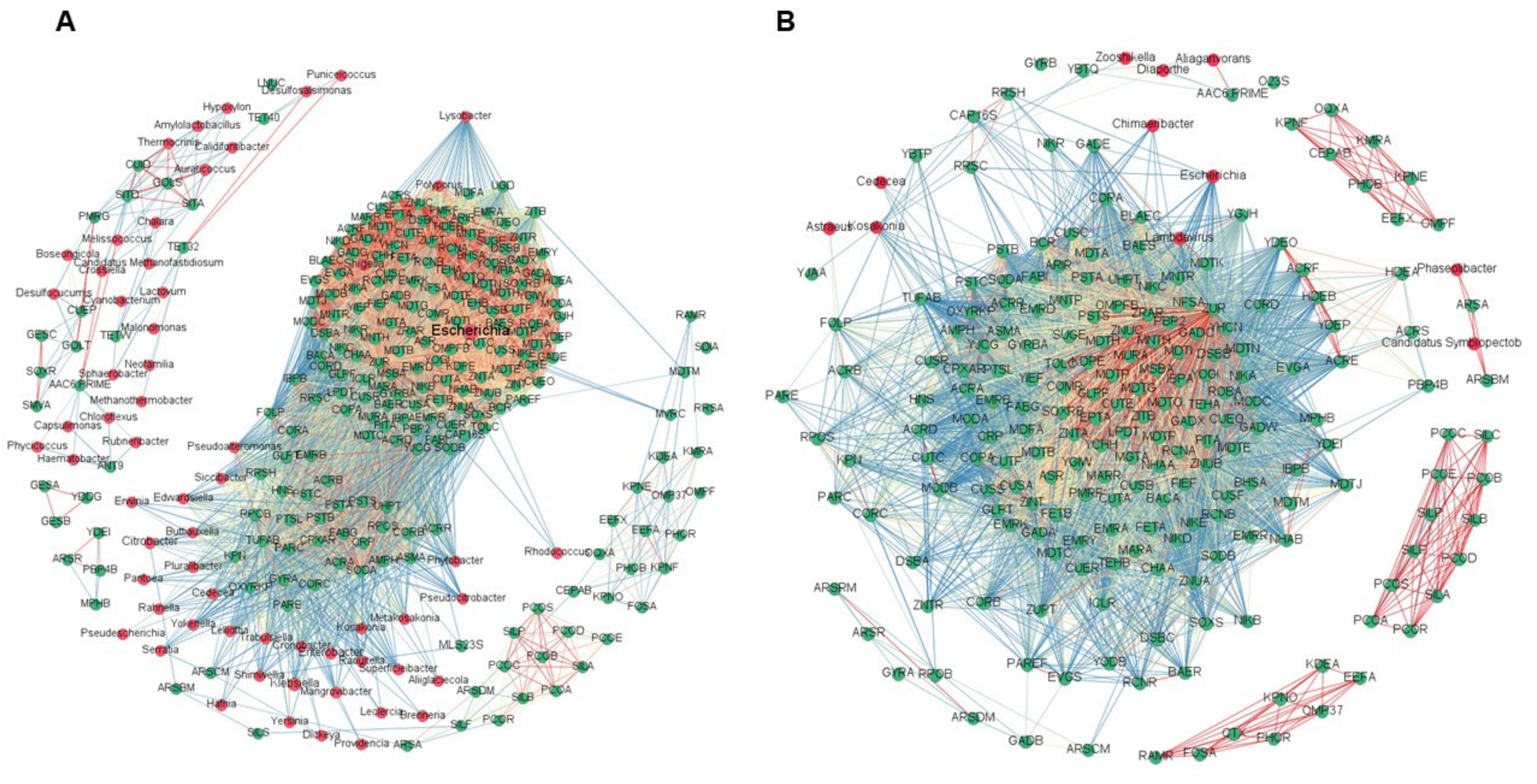


**Additional file 20. ARG-ARG and ARG-taxa connections are different for individuals infected with and recovering from *Campylobacter*.** Correlation co-occurrence networks were constructed in Gephi 0.9.2 using Spearman's Rank correlation coefficients generated with the R-package 'Hmisc' (v4.5-0) for cases infected with *Campylobacter* (A) and recovering follow-ups (B). These networks display all ARG-ARG and ARG-taxa connections; taxa-taxa connections were excluded for clarity. Correlations included in the network all passed a cutoff of $\rho>0.80$ and q-value $< 0.05$. Nodes are colored by their identity as a taxonomic genus (red) or ARG group (green). Nodes are sized based on their overall abundance among samples; the larger the node, the more abundant. Nodes with $\geq$ 2 connections were included (i.e. degree cutoff=2). The edge color displays the strength of correlation, with blue demonstrating relatively weaker correlations (yet still >0.80), yellow showing medium correlation, and red showing strong correlation. Nodes are labeled with their corresponding genus or ARG group.
